# Cadherin-11 dimerization multi-site kinetics: combined partial unfolding and strand-swapping

**DOI:** 10.1101/2020.12.21.423864

**Authors:** Hans Koss, Barry Honig, Lawrence Shapiro, Arthur G Palmer

**Affiliations:** Department of Biochemistry and Molecular Biophysics, Columbia University Irving Medical Center, 701 West 168^th^ Street, New York, NY 10032, USA; Zuckerman Brain Mind Behavior Institute, Columbia University, 3227 Broadway, New York, NY 10027, USA; Department of Systems Biology, Columbia University Irving Medical Center, 1130 St. Nicholas Avenue, New York, NY, 10032, USA; Department of Medicine, 630 West 168^th^ Street, Columbia University Irving Medical Center, New York, NY, 10032, USA

**Author notes:** Address correspondence to A.G.P.

## Abstract

Cadherin extracellular domain 1 (EC1) mediates homophilic dimerization in adherens junctions. Conserved Trp2 and Trp4 residues in type II cadherins anchor the EC1 A-strand intermolecularly in strand-swapped dimers. Herein, NMR spectroscopy is used to elucidate the roles of Trp2 and Trp4 in Cadherin-11 dimerization. The monomeric state, with the A-strand and Trp side chains packed intramolecularly, is in equilibrium with sparsely populated partially and fully A-strand-exposed states, in which Trp2 (and Trp4, respectively) side-chain packing is disrupted. Exchange kinetics between the major state and the partially (fully) A-strand-exposed state is slow-intermediate (intermediate-fast). A separate very fast process exchanges ordered and random-coil BC loop conformations with populations dependent on A-strand exposure and dimerization status. Additionally, very slow processes connect the folded A-strand-exposed conformation to partially unfolded states, which may represent additional domain-swapping intermediates. The dimerization mechanism of type II cadherins is revealed as coupled folding and strand-swapping.

## Introduction

Classical cadherins, which include the type I and type II families in vertebrates, are calcium-dependent cell-cell adhesion proteins (Brasch et al., 2012). Type II cadherin ectodomains contain five “extracellular cadherin” (EC) domains. The membrane-distal cadherin EC1 domains from cadherin molecules on the surfaces of opposing cells form dimers in the process of cell-cell adhesion. The greatly varying dimerization propensities of various cadherin homo- and heterodimers is thought to play an important role in tissue patterning (Brasch et al., 2018). EC1 is the primary determinant of type II cadherin dimerization specificity; the high level of similarity of dimeric interfaces in type II Cadherins raises the question of how this specificity is achieved (Patel et al., 2006). While isolated EC1 constructs can form stable dimers, and all intermolecular contacts are located between EC1 domains, binding of Ca^2+^ rigidifies the connections between successive EC domains. Dimerization of EC1-EC2 protein constructs is accelerated by formation an “X-dimer” kinetic intermediate (Harrison et al., 2010, Li et al., 2013).

Type II cadherin EC1 domains form strand-swapped dimers, with the N-terminal A-strand undocking from an intramolecular site in the monomer to bind an identical intermolecular site in the partner protomer (Shapiro et al., 1995, Patel et al., 2006). The side chains of residues Trp2 and Trp4 near the N-terminus are packed intramolecularly in the monomer conformation and intermolecularly to anchor the A strand in the domain-swapped conformation (Patel et al., 2006). Prior work has suggested that Cadherin 8, a type II Cadherin, can access an A-strand-exposed conformation in the monomer state to facilitate dimerization (Miloushev et al., 2008). However, the existence of an A-strand-exposed intermediate in Cadherin dimerization is still elusive and the precise role of Trp4 in type II Cadherins is unknown (Sivasankar et al., 2009, Miloushev et al., 2008).

Partially or fully unfolded intermediates have been described for domain-swapping dimerization processes, compatible with the concept that domain swapping is intermolecular protein folding (Moschen and Tollinger, 2014, Rousseau et al., 2003). Partial unfolded intermediates have been detected in β-strand swapping of a GBl mutant (Byeon et al., 2004) and in (-terminal domain-swapping of RNAse A (Esposito and Daggett, 2005, Liu et al., 2001). Domain swapping also can require full unfolding, as exemplified for Cyanovirin-N protein (Liu et al., 2012). Unfolding can promote domain swapping in a variety of ways. (1) High concentrations may favor non-thermal ‘melting’ of the protein in which a protein is acting as a solvent (Daoud et al., 1975, Gronenborn, 2009). Large hydrophobic surfaces could promote the formation of encounter complexes, as they occur in induced-fit mechanisms (Spadaccini et al., 2014). Accordingly, the presence of monomeric and dimeric intermediates of M^pro^-C of SARS-CoV main protease in which the C-terminal A5-helix is unfolded may provide a hydrophobic environment to enable a1-helix strand-swapping (Kang et al., 2012, Liu and Huang, 2013). (2) Simulations of the strand-swapping SH3 domain of epidermal growth factor receptor pathway substrate 8 (Eps8) show that unfolded intermediate states increase competition between intra- and inter-chain interactions by virtue of their encounter complex interfaces (Yang et al., 2004). (3) Unfolded states can serve as free energy traps in the domain-swapping process while avoiding an overly rugged energy landscape during dimerization (Yang et al., 2004).

The present work characterizes the conformational states of the monomeric form of the Type II cadherin-11 EC1 (Cad11-EC1) domain(Patel et al., 2006, Chang et al., 2010, Sfikakis et al., 2017). Extensive NMR spectroscopic measurements, including chemical shift perturbation and relaxation dispersion, were conducted on wild-type (WT) and designed mutants of Cad11-EC1, including dependence on temperature and pressure. The results describe an ensemble of folded, locally disordered, A-strand exposed, and partially unfolded conformational states of Cad11-EC1 in solution. This landscape of conformations interconverting at different time scales reveals dimerization of Cadherin Type II domains as coupled folding and A-strand swapping and further emphasizes the importance of conformational plasticity of the Trp2 binding pocket and the destabilizing influence of Trp4 in the mechanism.

## Results

The key NMR probes of the conformational dynamics of the Cadherin-11-EC1 (WT-C11) domain are Trp2 and Trp4 in the A-strand of the domain, together with the flanking BC (residues 29-34), DE (residues 52-55), and FG (residues 80-84) loops, as shown in ***Fig. 1a***.

**Fig. 1:**
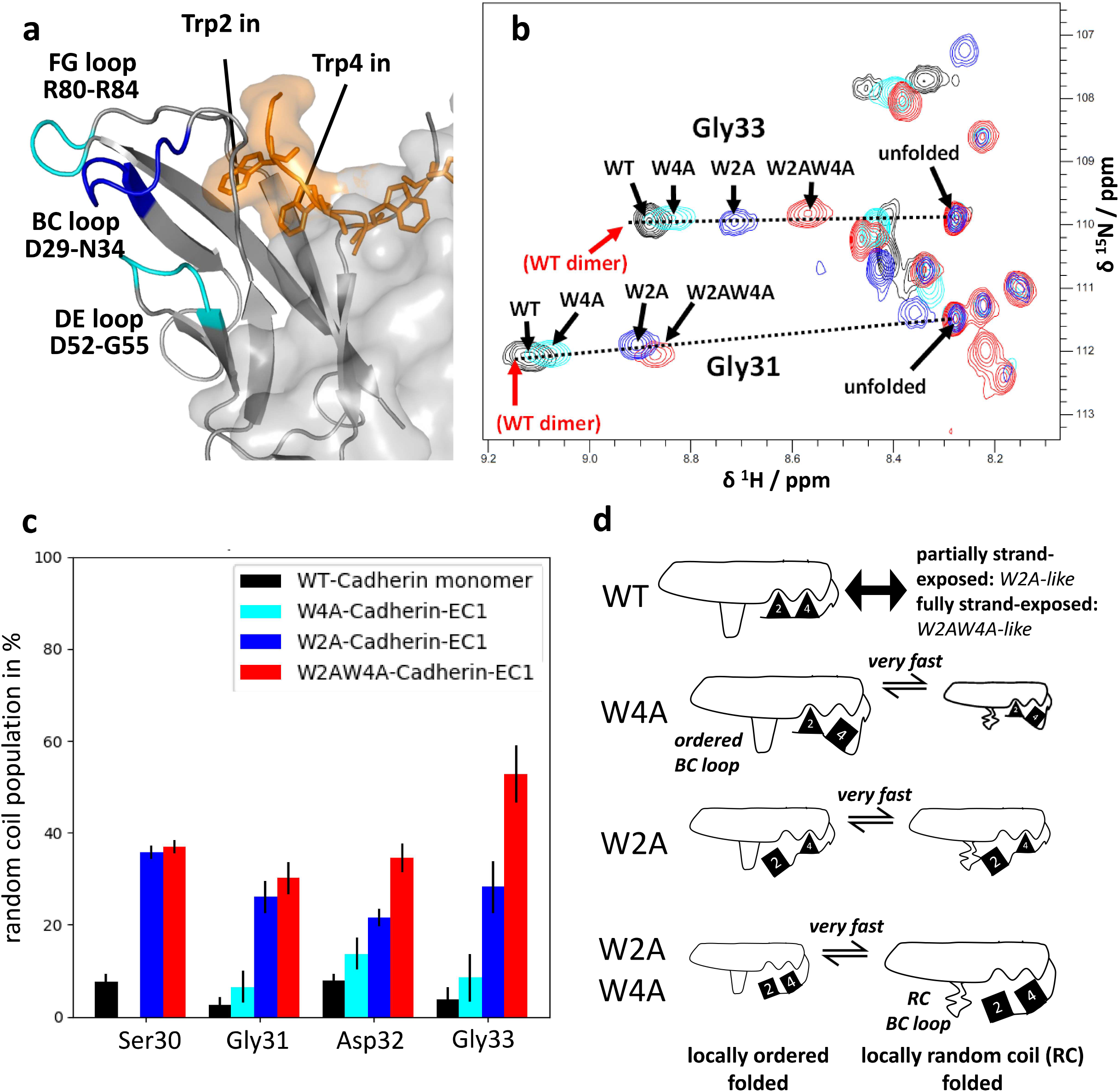
Chemical exchange in Trp2 and Trp4 mutants. **a: Structural features of Cad-11 EC1.** Orange : N-terminal strand with Trp2 Trp4 side chains of homodimerized Cad-11 EC1 inserted in the intermolecular binding pocket. The BC, DE, and FG loops, which display unique dynamic features without confounding secondary chemical shift perturbations, are indicated. **b: Superposed** ^**1**^**H**,^**15**^**N-HSQC spectra for WT, W2A, W4A, and W2AW4A mutants of Cadherin-11-EC1**. The position of the WT dimer peaks (not visible in this dilute sample) is indicated. Dotted lines are drawn from the dimer resonance to the RC resonance for each amino acid residue. The random coil (RC) peak position is common to all constructs, but intensities vary greatly between mutants. **c: Semi-quantitative estimation of RC populations from projections on the** ^**1**^**H**,^**15**^**N-vector**. The end points for the RC population destination are the ordered dimer and RC states. The RC population for different constructs varies between residues (see also *Fig. S4*). **d: Very fast exchange based on the mutant spectra**. Sizes of cartoons represent populations (not to scale). Positions and status of Trp2, Trp4, and BC loop are indicated.

### Monomer and dimer NMR spectra

The ^1^H-^15^ N correlation spectrum of Cad11-EC1 ***(Fig. 3a)*** displays substantial line broadening and, in many cases, more than one resonance peak per residue. By comparing spectra at different concentrations, we were able to assign resonance peaks to a main monomer state, a main dimer state, or an overlap of each. Assignments of resonances to monomer or dimer states were confirmed by comparing experimentally obtained relaxation rate constant ratios with those calculated from the crystal structure ***(Fig. S1*)**. A total of 44 residues (47 % excluding prolines) were assigned for the main monomer state of WT Cad11-EC1 ***(Table S1*)**. The two-domain Cadherin-11-EC1-EC2 protein construct is difficult to access under conditions suitable for NMR spectroscopy. However, the overlap between ^1^H-^15^N-HSQC spectra for Cad11-EC1 and Cadherin-11-EC1-EC2 indicates that the properties of Cad11-EC1 are similar in the two-domain context ***(Fig. S2*)**.

### Trp2 and Trp4 mutants reveal very fast dynamics between ordered and random coil states of the BC loop

We introduced W2A and/or W4A mutants to test the role of individual Trp (W) residues. New resonance peaks were observed in NMR spectra of the mutant domains (assignments and intensities in ***Table S2*)**. A subset of the new resonance peaks corresponds to the random coil (RC) states : the correlation between predicted random coil ^15^N-shifts and assigned ^15^ N resonances is 96% ***(Figs. S3a-S3b)***. The ^1^H,^15^ N-HSQC peaks for BC loop residues Ser30-Gly33 in wild-type and mutant domains can be placed on a line connecting the peaks for WT-dimer and RC monomer ***(Figs. 1b, S3c-S3e)***. This correlation indicates that a structurally ***ordered*** state kinetically exchanges with a disordered ***random coil*** (RC) state. The resonance peaks for dimer states are shifted in the opposite direction from the RC states. Thus, the fraction of RC state (equal to the degree of ***local unfolding)*** in the BC loop generally follows the order: WT(dimer)-WT - W4A - W2A-W2AW4A - RC, with WT-dimer being predominantly ordered (states without a designation as dimer are monomeric).

The RC state population varies for each mutant ***(Fig. 1c)***, suggesting that the fast exchange equilibria are partially independent. The RC population of the W2AW4A mutant obtained from ^13^C_α_ resonance peak positions for residues Ser30, Asp32 and Gly33 match the results from the analysis of ^1^H-^15^N spectra ***(Fig.S3f)***.

Resonance peak broadening due to this very fast exchange process could not be suppressed in high power relaxation dispersion experiments *(vide infra);* therefore, the exchange process is very fast on the NMR chemical shift time scale (< 10 - 100 µs). The peak patterns observed for residues in the DE and FG loop, particularly Asp81 and Asn83, in WT and mutant C11 could suggest a very fast exchange kinetic process as well; however, the linear correlation of the chemical shifts for the wild-type and mutant proteins does not extend to the RC peak positions. Alternatively, the peak patterns in the DE and FG loops could be a consequence of local chemical shift perturbations by the structural changes in the neighboring BC loop.

### Population and structure of partially unfolded states depend on Trp status

The Cad11 –EC1 monomer structures associated with RC resonance frequencies are ***partially unfolded states (Figure 2)***. The resonance peaks for folded states are resolved from the RC peaks and hence the folded and partially folded states exchange slowly on the NMR chemical shift time scale. The overall intensities of RC peaks for the W2A mutant are 4.3 times smaller than for the W2AW4A mutant when scaled to the average folded state intensity, showing increased population the partially folded state for the W2AW4A mutant ***(Fig. 2a)***. RC peak intensities for the mutant proteins are greatly influenced by the populations of these states and by the effects of the local structural context on effective rotational correlation times. Therefore, we normalized RC peak intensities by the average RC intensity of all assigned residues up until and including DE loop (residues 1-55). This normalization yielded identical residue-wise intensity profiles for large sections of W2A-Cl l and W2AW4A-C11 ***(Fig. 2b, 2c*)**, indicating that large regions of partially unfolded states of W2A and W2AW4A mutants are structurally identical, pointing to a common unfolding pathway. The RC peak intensities from Val78 to lle93, including the FG loop and the only assigned G strand residue, generally increase in the order W2A mutant< W4A mutant< W2AW4A mutant. Projection of these increased intensities on the structure suggests that disruptions of interactions between Trp4 and G strand residues are responsible for increased local unfolding ***(Fig. 2c*)**. Arg80 is an outlier in terms of relative intensity differences between mutants ***(Fig. 2b*);** inspection of the loop region reveals that its interactions with the BC and the DE loop are stabilized upon FG loop unfolding ***(Fig. 2d*)**.

**Fig. 2:**
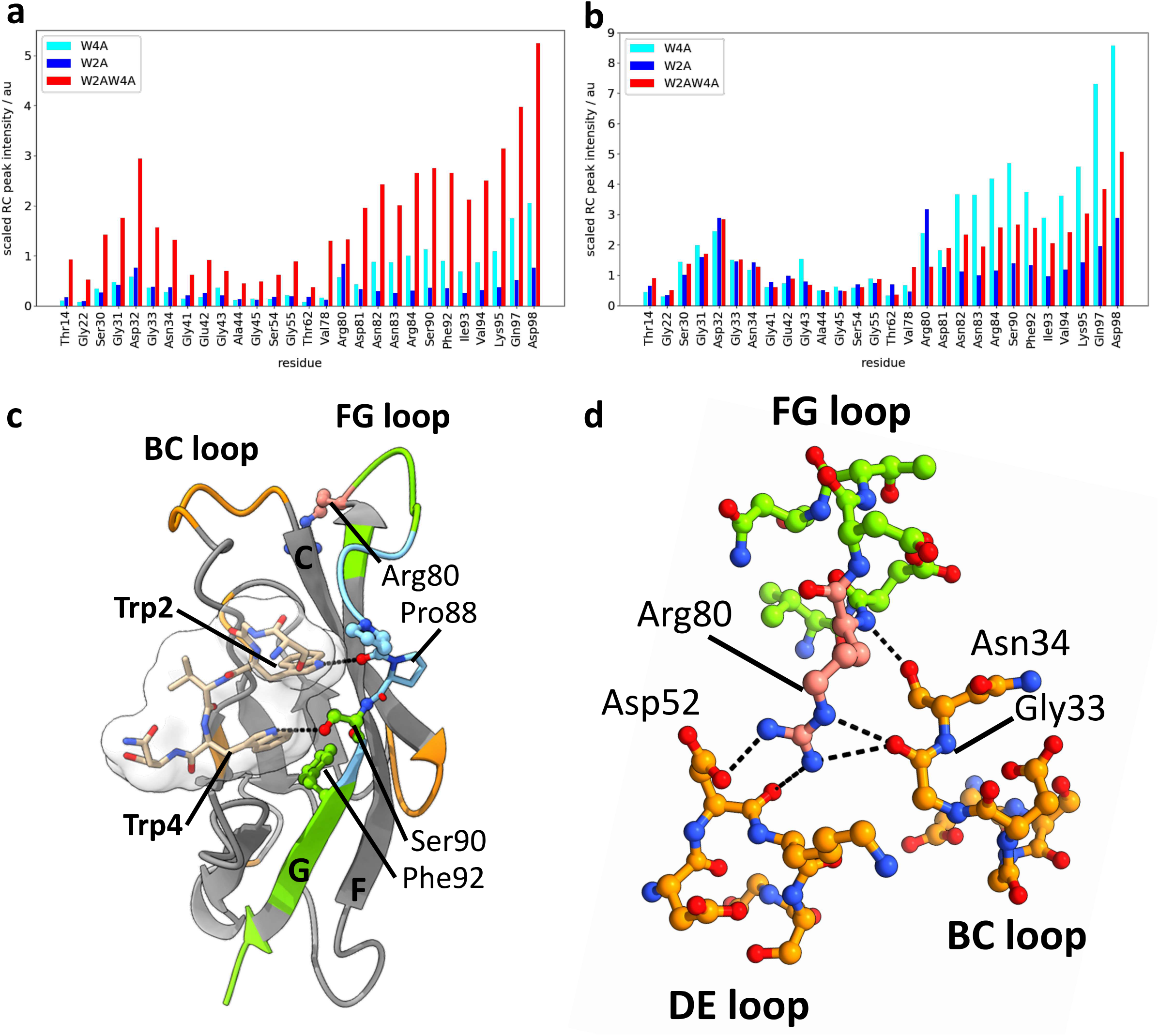
Partially unfolded states in in Trp2 and Trp4 mutants. **a. Partially unfolded populations in the Trp mutants.** Intensities were scaled to the average intensity of folded peaks. The population of the partially unfolded state is highest in the W2AW4A mutant. **b. Intensity data give clues about structural similarity of partially unfolded states**. Intensities were scaled to the average intensity of unfolded peaks for all residues in sequence up until residue 55. The data suggest similar partially unfolded structures except for the region encompassing the FG loop and G loop residue 193 which are more destabilized in W2AW4A and W4A. Arg80 shows a different pattern, which is rationalized in panel d. **c: Destabilization of the FG loop and the G strand in the W2AW4A mutant**. Orange: Partial unfolding which is identical for the W2A and the W2AW4A mutant. Green: Residues for which the W2AW4A mutation leads to more unfolding. Cyan: G strand residues linking residues colored in green; the marked residues are highly conserved only in type II Cadherins and connect structurally to Trp2 and Trp4. **e. Structural connections between loops**. Residue 80 uniquely connects the FG, BC and DE loops, commensurate with its stabilization upon unfolding of the FG loop.

**Fig. 3:**
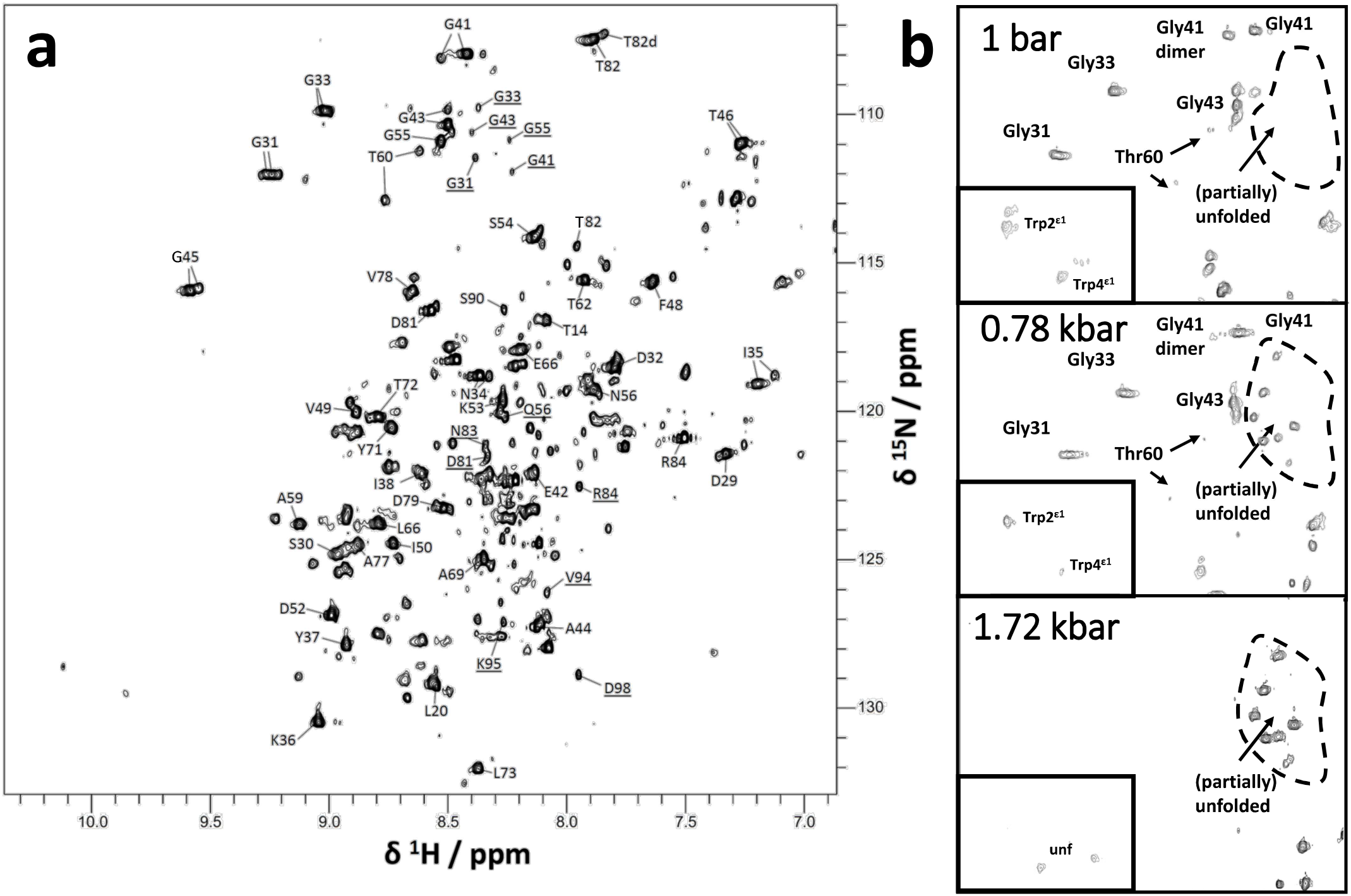
^1^H,^15^N-TROSY-HSQC spectra of Cad11-EC1-WT at different pressures. **a:** ^**1**^**H-**^**15**^**N correlation spectra recorded at 900 MHz, 160 mM, and 298 K, and 1 bar pressure**. Duplicate assignments mostly refer to monomer/dimer pairs. Underlined peaks refer to random coil peaks, which are hardly detectable in WT samples. The two marked regions refer to the spectral regions in the right panel. **b: Pressure series recorded at 800 MHz, 160 µM and 285 K**. The pressures are shown in the figure. The corresponding areas in the full ^1^H,^15^N spectrum are marked in the left panel; see also *Fig. S4*.

All G strand residues interacting with Trp2 and Trp4 are fully conserved in type II, but not type I Cadherin, commensurate with partial unfolding as an important and specific feature of type II Cadherin dimerization kinetics via interactions of both Trp2 and Trp4 with the G strand.

### High pressure NMR experiments

The ^1^H,^15^ N-HSQC spectra obtained from static high-pressure titrations from 1 bar to 2.07 kbar show dramatic changes in numbers, intensities, and frequencies of resonance peaks ***(Figs. 3, S4*)**. Pressure-dependent intensities of RC and folded peaks reveal unfolding under pressure. Increases in folded monomer peak intensities at low pressures arise in part from pressure-dependent dimer-monomer transitions; however, local packing effects or chemical exchange processes further modulate site-specific peak int ensit ies. Similar to recent work on Enzyme I, we use high-pressure experiments to monitor order-disorder transitions of loops functionally linked to dimerization (Nguyen et al., 2021).

### Multi-step pressure unfolding reveals intermediate- and high-pressure partially unfolded states

At 1.72 kbar, the ^1^H,^15^ N-HSQC spectrum is consistent with a mostly unfolded state of WT Cad11-EC1 ***(Figs. 3b, S4*)**, in which the resonance frequencies for all assigned residues agree with RC values. Correspondingly, ∼20 new pressure-dependent resonance peaks match the assignments of the RC peaks for the W2AW4A mutant.

Pressure-dependent resonance peak intensities and frequencies were fit with a generic spline function that depends on two linear features and a critical pressure at which the linear slopes change ***(STAR Methods)***. The behavior of the intensities and resonance positions of RC peaks suggests that unfolding from 0 kbar to 1.72 kbar occurs in at least two distinct stages, from 0-0.97 kbar and 0.97-1.72 kbar, thus defining ***intermediate-pressure* (IP-UNF)** and ***high-pressure partially unfolded* (HP-UNF)** states ***(Figs. 3, 4 a-c, S4-S7*)**. Resonance peaks for the IP-UNF partially unfolded state shift uniformly towards the resonance positions for HP-UNF at pressures > 0.97 kbar, commensurate with fast exchange between these states.

**Fig. 4:**
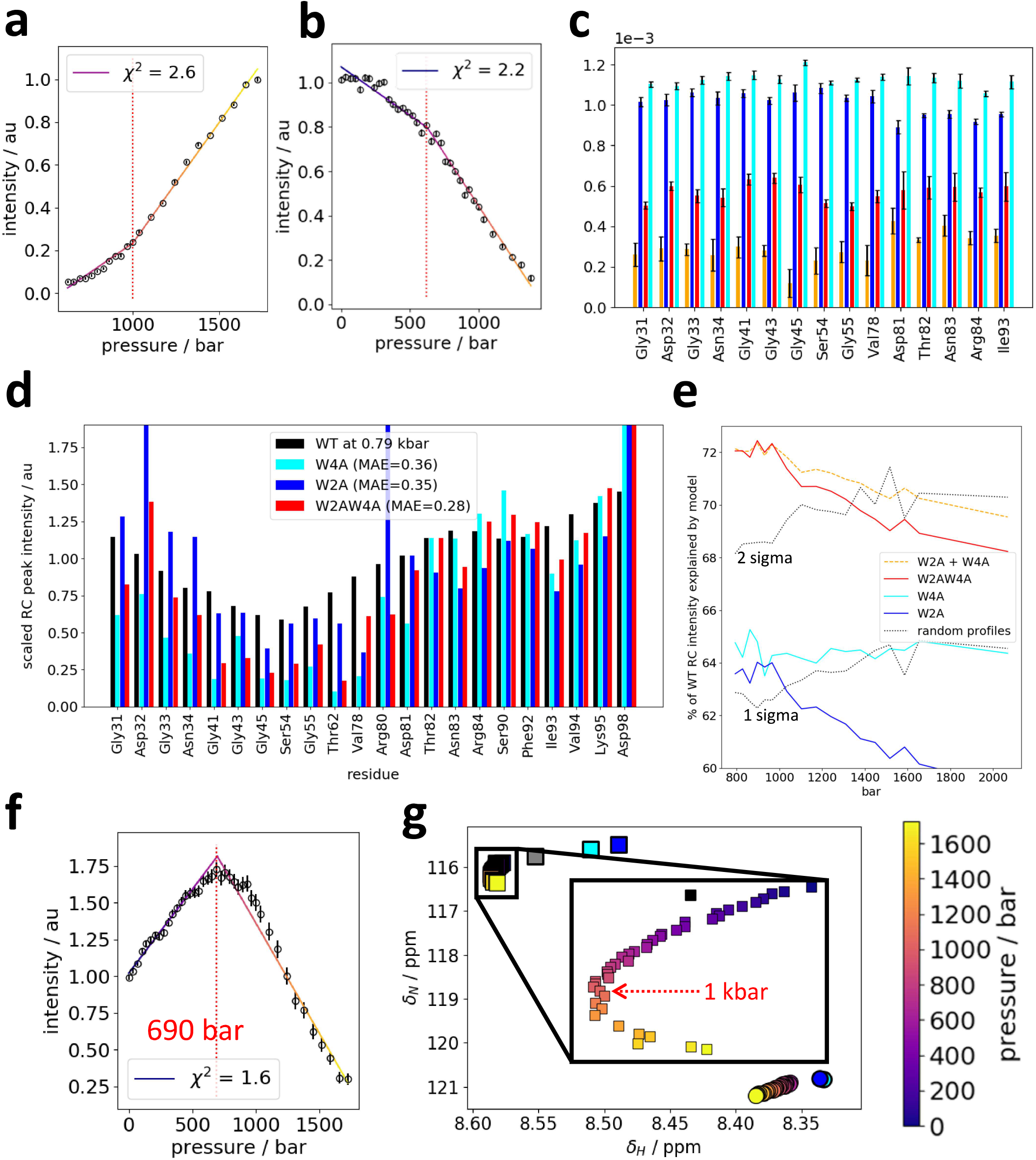
Analysis of pressure dependence of peak intensities. **a,b: Pressure dependency of Gly33 peak intensities at 285K**. A first-order spline with three points (two straight lines) was used to characterize each of the three intensity profiles; panel a shows the intensity of the RC (unfolded) peak, panel b the intensity of the folded peak. The dotted red line indicates the pressure of the central critical pressure for the three-point first order spline fit. For more examples see *Fig. S5. χ*^2^ is shown in the figure legend. **c: Fitted slopes for intensity dependency on pressure of unfolded peaks at 283 K and 298 K**. Three-point first-order spline (right panel) fits give two separate slopes: The low-pressure slope for 283K (red) and 298K (orange), and the high-pressure slopes for 283K (cyan) and 298K (blue). **d**,**e: Fitting Trp mutant RC intensity profiles to profiles of high-pressure WT states**. Of the three Trp mutant profiles, the W2AW4A mutant intensity profile is best to be scaled to match the normalized WT intensity profile at 0.79 bar; the color assignment and mean average error (MAE) is shown in the legend (panel d). The share of the pressure-dependent WR WT intensity which can be explained (1 - MAE) by various mutant intensity profiles is shown in panel e; the upper sigma and 2 sigma boundaries of the results for fits with random intensity profiles are shown. **f**,**g: Pressure dependency of folded Asp81 peak intensities (panel f) and chemical shifts (panel g) at 285K**. The intensities in panel f were fitted in the same manner as described in panels a and b. The color of the calculated curve indicates pressure. The same pressure scale is used for intensities in panel g, alongside a color bar (right) translating colors into pressures. Other symbols in panel g: squares - folded peaks; circles - unfolded (RC) peaks; blue: W2AW4A mutant peak; cyan: W2A mutant peak; grey: W4A mutant peak: black: WT peak for comparison with mutants. The critical pressures for intensities (panel f) and chemical shifts (panel g) differ, indicating a two-step unfolding process: from 690 bar, general decline of the population (slow exchange type); then from around 1 kbar, local unfolding. Resonance frequency shifts in ^1^H,^15^N-TROSY spectra recorded at pressures from 1 bar to 1.72 kbar result from population changes between conformational states in fast exchange. The folded peak for Asp81 shifts away from the unfolded peak at low pressures and “changes direction” to move towards the unfolded state only above ≈ 1 kbar (inset).

Fitting RC intensity profiles for mutant constructs to the pressure-dependent WT RC intensity profile reveals that the partially unfolded W2AW4A state explains 72% (mean average error, MAE= 0.28) of the normalized intensity profiles at 0.79 - 0.97 kpsi ***(Figs. 4d,e)***. This is significantly more than for random intensities (2 sigma interval : 41-68 % at 0.79 bar) and W4A or W2A (both 64 %). A linear combination of W4A and W2A explains 72 % of the intensit ies. Comparing the scaled intensity profiles suggests that the W2AW4A intensity profile generally matches the FG loop and G strand region better than W2A at pressures up until 0.97 kbar ***(Fig. 4d)***. We conclude that the partially unfolded intermediate-pressure states are similar to the partially unfolded W2AW4A state or a combination of partially unfolded A-strand-exposed states.

### Pressure-dependence of RC intensities and of folded peak chemical shifts indicates locally varying stability

Pressure-dependent shifts in the ^1^H and ^15^N resonance frequencies for peaks associated with folded conformations arise from shifts in the populations of alternative states that are in fast exchange on the NMR chemical shift timescale ***(Figs. 4g, S6-S8)***. Resonances for folded conformations of residues 31-34 in the BC loop peaks shift towards the HP-UNF peak positions as pressure increases, suggesting that the HP-UNF state is locally accessible for BC loop residues even at pressures < 0.97 kbar. In contrast, the peak positions for residue Ser54 in the DE loop and residues Asp81 and Asn83 in the FG loop shift non-linearly with increasing pressure and only towards the HP-UNF peak positions only at high pressures(⪆ 0.97 kbar, for Ser54 at 298K; ***Figs. 4g, S6-S8)***, notwithstanding an intensity decrease at ⪆ 0.69 bar due to a diminishing folded protein population ***(Fig. 4f)***. This result also suggests that the DE and FG loops are in very fast exchange between two ordered conformations, consistent with mutational results, and that they are more pressure-stable than the BC loop.

### Relaxation dispersion experiments for multi-site kinetics

Carr-Purcell-Meiboom-Gill (CPMG), CEST, Hahn-echo, and high power spin-lock ^15^N NMR experiments for WT Cad11-EC1 were used to characterize multi-site kinetics ***(Fig. 5)*** over a wide range of µs-ms timescales, while reducing ambiguity in the subsequent multi-site fitting time scales ***(STAR methods)*** (Palmer and Koss, 2019).

### Three-site model for strand exposure kinetics

Global three-site fits were performed for 12 residues ***(Figs. 5, S9-S10)***. We find slow-intermediate exchange (*k*_*ex*_ = 664 − 704 s^-1^) between a major (A-strand-bound) monomeric state (population: 96.77 -96.87 %) and a partially A-strand-exposed sparsely populated state (1.56 − 1.63 %) and intermediate-fast (*k*_*ex*_ = 2930−3217 s^-1^) exchange between the major state and a fully A-strand-exposed state (1.54−1.59 %), with a small concentration dependency of the partially exposed state population and the rate constants ***(Fig. 7, Tables S3-S5)***. The identified chemical shifts of the partially A-strand-exposed state map to those of the folded conformation of the W2A mutant, suggesting that this mutant is a model of the partially-strand exposed state of the WT domain. The shifts of the fully A-strand-exposed states mostly map to those of the folded conformation of the W2AW4A mutant with a slightly higher RC share, suggesting that the W2AW4A mutant is only mostly A-strand-exposed. The model is supported by qualitative analysis of ^1^H-CPMG relaxation data recorded for [U-^2^H, U-^15^N]-labeled samples ***(Fig. S11*)**. The data do not resolve whether direct exchange occurs between the partially and fully A-strand-exposed states. A model incorporating exchange between the partially and fully A-strand-exposed states is compatible with the same chemical shifts of the sparsely populated states, but yields different kinetic parameters: exchange between the A-strand-bound and the partially A-strand-exposed states would be *k*_*ex*_ = 266 - 334 s-^1^, and exchange between the sparsely populated states k_ex_ = 816 - 900s ^-1^ ***(Fig. S13a, Tables S3-S5)***.

**Fig. 5:**
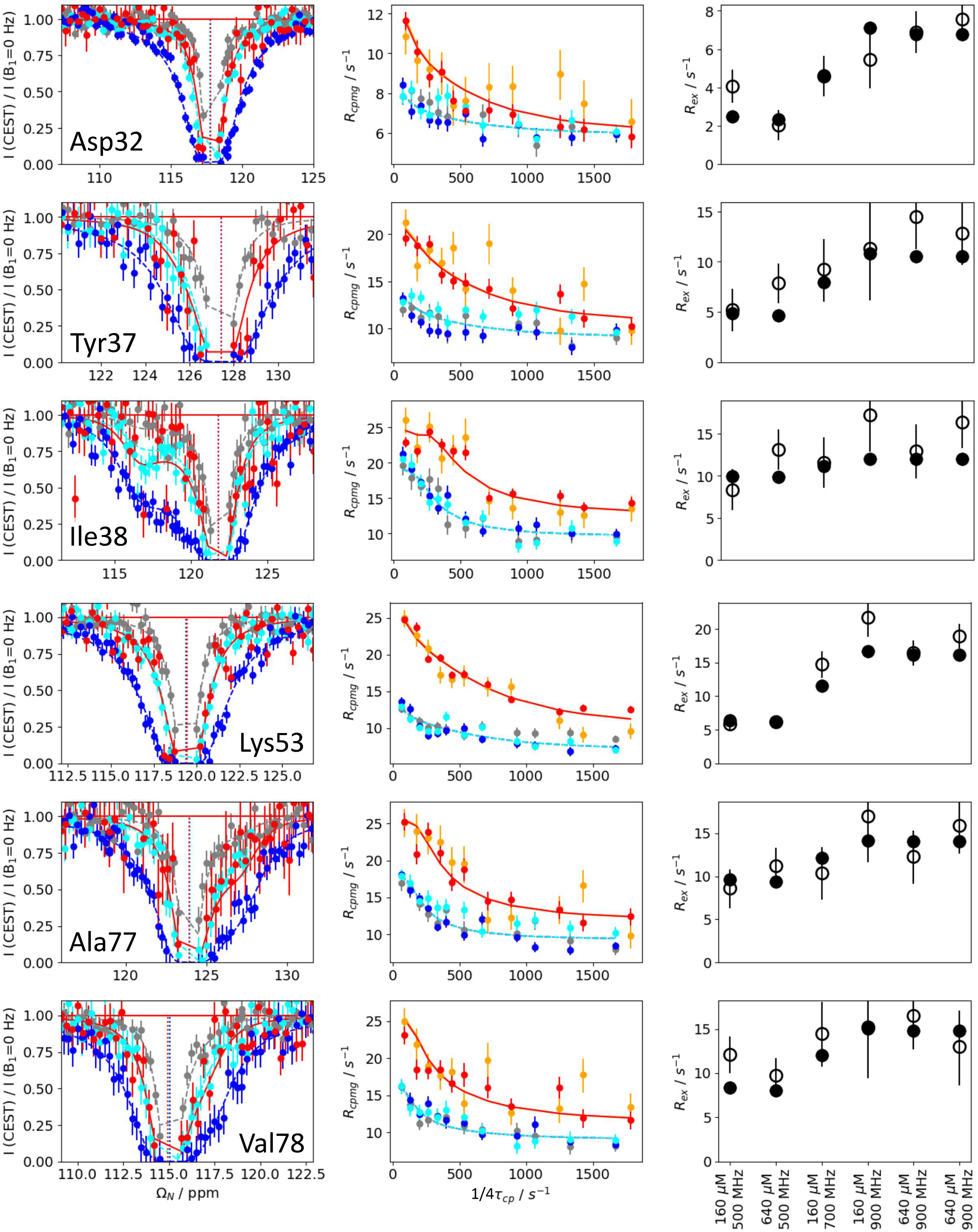
^15^N-relaxation dispersion reveals three-site exchange. Six of the 12 globally fitted residues have been selected to exemplify various type of chemical exchange observed within Cadherin-11-EC1 domain: Val38 : mostly slow-intermediate exchange; Asp32, Lys53: mostly fast-intermediate exchange; Tyr37, Ala77, Val78: combination of slow-intermediate and fast-intermediate exchange. Left panels: ^15^N-CEST at at c = 640 µM, B_0_ = 800 MHz, B_1_= 35 Hz (red) or at c = 160 µMand B_0_=700 MHz (grey- B_1_ = 15 Hz, cyan - B_1_ = 15 Hz, blue - B_1_ = 50 Hz); center panels: ^15^N-CPMG at c = 160 µM (grey, orange), c = 640 µM (blue, cyan, red), B_0_= 500 MHz (grey, cyan, blue), B_0_ = 900 MHz (orange, red); right panels: ^15^N-high power spin-lock experiments for *R*_*ex*_ determination, with conditions listed along the x axis with experimental (open circles) and calculated (filled circles) values. For the global fit, ^15^N-CEST data were converted to *R*_1ρ_-type data as described elsewhere (Palmer and Koss, 2019), using *R*_1ρ_ multi-site approximations for initial calculations (Koss et al., 2017). CPMG data were fitted with an the exact solution or an approximation (see *STAR Methods)* for the triangular state (Koss et al., 2018). For more data, see *Fig. S9*.

### Concentration dependence of fast-intermediate exchange between A-strand-bound and fully A-strand-exposed

Many chemical shifts in ^1^H-^15^N correlation spectra for Cad11-EC1 display a strong Bo field dependence ***(Fig. S12a)***, which itself is a signature of chemical exchange. The B_0_-induced chemical shift perturbations between 500 and 900 MHz (Δδ_B0_) correlate linearly between low (160 mM, Δδ_B0,low_) and high (640 mM, Δδ_B0,high_) concentration samples, ruling out most arti facts, systemic and sample-related errors contributing to the shift perturbations ***(Figs. S12c, S12d)***. The average B_0_-dependent shift perturbation is 10% greater at high concentration, suggesting that the underlying chemical exchange process has a degree of concentration dependence. The B_0_-dependent ^15^N chemical shift differences at high concentration also are correlated (R = 0.72) with the ^15^N chemical shift differences between the major conformation and fully-strand exposed conformation detected in relaxation dispersion experiments ***(Fig. S12b)***.

### Temperature dependence reveals features of partially unfolded states

We recorded relaxation ^15^N-CPM G dispersion at 285 K and 298 K. The slow-intermediate process between WT and the partially A-strand-exposed (W2A-like) state is temperature independent. The slow-intermediate process was mapped to the structure by directly comparing the peak intensities in ^1^H,^15^N-HSQC spectra recorded at different fields and temperatures ***(Fig. 6*);** no chemical shifts or data obtained from relaxation measurements were needed to perform this procedure. We find that the slow-intermediate process indeed maps to the W2A binding pocket. The line-broadening arising from exchange between WT and the fully A-strand-exposed state (W2W4-like) decreases with increasing temperature, as expected for an intermediate-fast time scale kinetic process. As already noted, the fully A-strand-exposed state is closely identified with the folded W2AW4A-like mutant and the WT IP-UNF states.

**Fig 6:**
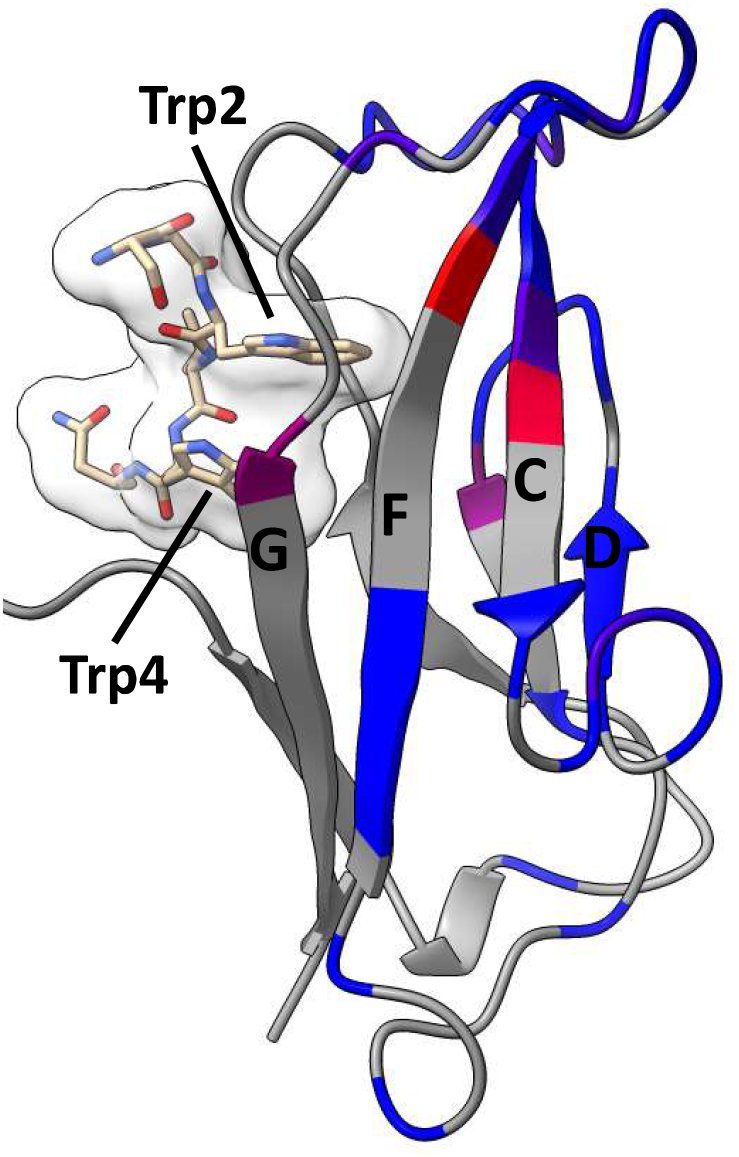
Structural characteristics of the sparsely populated partially strand exposed state. This state maps to the Trp2 binding pocket, in line with Trp2 exposure. The peak intensity ratio I^298K,500MHz^ *I ^285K,900MHZ^/I^298K, 900MHz^*I ^285K, 500MHz^ is projected on the A-strand-bound structure (blue-violet - red). The result confirms the expectation from relaxation dispersion analysis that the ratio encodes the chemical shift perturbation between WT and the minor W2A-like state.

**Fig. 7:**
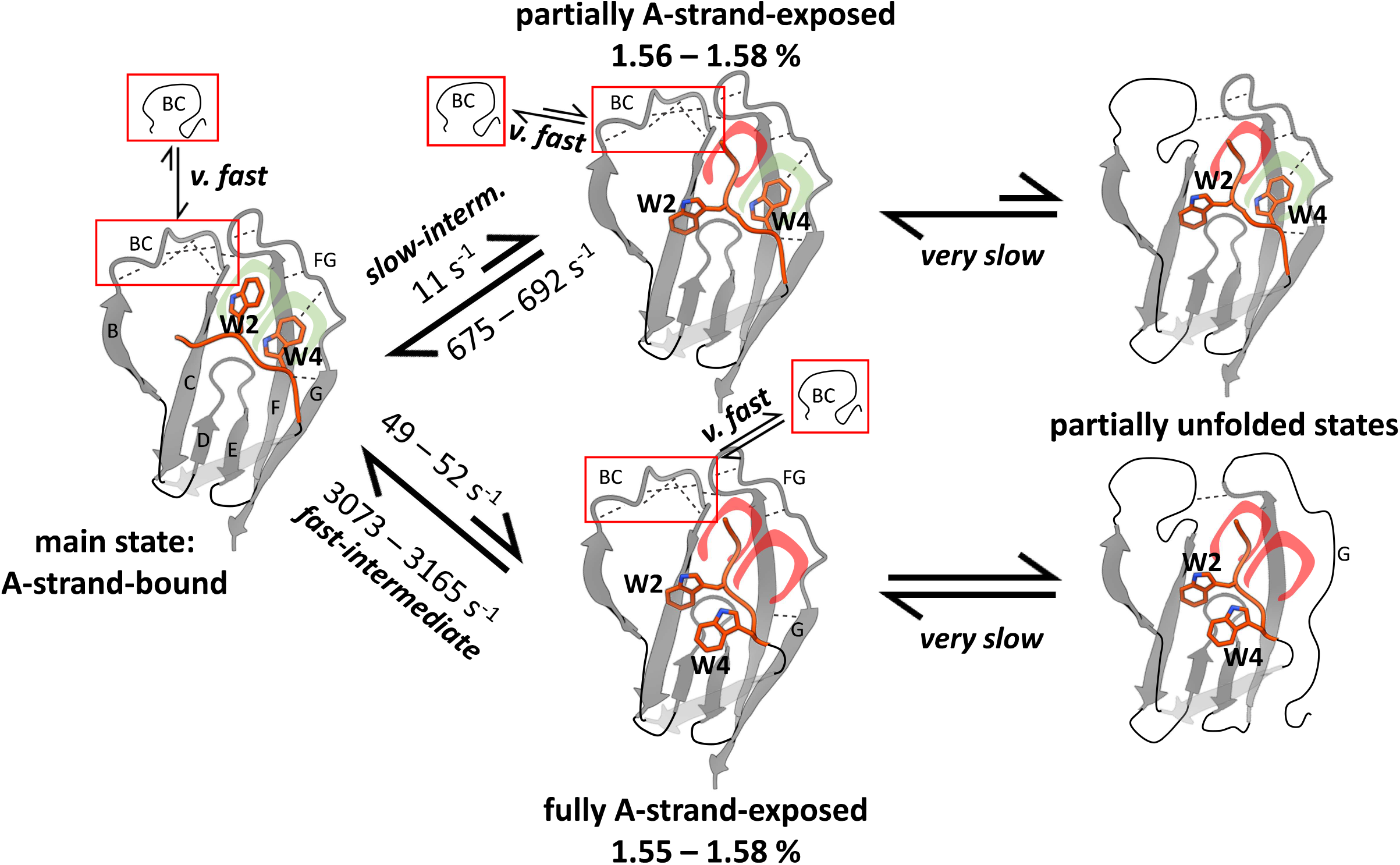
Chemical exchange in Cad11-EC1 WT. Scheme illustrating chemical exchange at various dynamic levels. The local very fast equilibria for BC loop are shown in red insets, with the equilibrium arrow indicating general propensity towards the ordered or the RC state. Exemplary contacts in the locally ordered states are shown as dotted lines, RC sections are shown as black solid lines. The binding status of the Trp2 and Trp4 pockets are illustrated in red (not bound) and green (bound). Important strand and loop labels are indicated in the schematic, simplified models. Slow-intermediate and fast-intermediate exchange have been established by relaxation dispersion. Resulting rate constant and population ranges for c = 640 µM are shown. Populations are normalized to the total population of folded states, excluding partially unfolded states. Significantly different results for c = 160 µM : p(strand-exposed) = 1.62 − 1.63 %; *k*_*31*_(partially A-strand-exposed → A-strand-bound) = 653 − 663 s^-1^; *k*_*21*_(fully A-strand-exposed → A-strand-bound): 2884 − 3059 s^-1^. For details and results including exchange between the A-strand-exposed states see *Fig. S13* and *Table S2*.

## Discussion

As a type II Cadherin, Cadherin-11 carries Trp2 and Trp4 as two key residues involved in the mechanism of strand swapping and dimerization. To investigate the roles of these Trp residues, we quantitatively characterized fast-intermediate timescale chemical exchange of WT-Cad11 between the main A-strand-bound and a sparsely populated fully A-strand-exposed (W2AW4A mutant-like) state, and slow-intermediate timescale chemical exchange between the main A-strand-bound and a sparsely populated partially A-strand-exposed (W2A mutant-like) state ***(Figs. 7, S13a*)**. The fully stand-exposed state also has both Trp2 and Trp4 unpacked from the intramolecular binding pocket, while the partially exposed strand has only Trp2 unpacked. Additional very fast exchange occurs between ordered and disordered (random-coil like) conformations of the BC loop, while the neighboring DE and FG loops are either in fast exchange between two alternative ordered conformations or structurally influenced by the unfolding process of the BC loop. The populations of the RC states of loops increase in the order dimer< monomer A-strand-bound < WT-partially A-strand-exposed < fully A-strand-exposed. Because exchange between A-strand-bound and partially A-strand-exposed states is slower than between A-strand-bound and fully A-strand-exposed states, we conclude that Trp2 exposure is not a required precursor to Trp2 and Trp4 exposure. Rat her, Trp2 and Trp4 detach from the core either simultaneously or via a Trp4-exposed state.

Mutation studies suggest that unbinding Trp2 or combined Trp2/Trp4 unbinding increases RC populations in BC loop. However, the A-strand-bound dimer state leads to a larger ordered population than the A-strand-bound monomer state and hence reflects stabilization of the dimer state compared to the monomer state. This implies that the maximum ordered population of the dimer is not merely a result of stabilization by the A-strand-bound state. This stabilization of the ordered state of BC loop in the dimer, but not in the monomer, might be a factor, in addition to conformational strain differences in the A strand (Vendome et al., 2011), to rationalize the preference for strand-swapping over self-insertion of the strand.

Very slow chemical exchange of folded states with partially unfolded states also depends on strand conformation and packing of Trp2 and Trp4. The effects of Trp2 and Trp4 exposure distinctly destabilize different sections of the EC1 domain. Thus, FG loop and G strand residues are destabilized more by combined exposure of Trp2 and Trp4 than by exposure of Trp2 alone.

The model of direct kinetic connections between folded and partially unfolded A-strand-exposed states is supported by three lines of evidence. (1) Trp mutant spectra, corresponding to A-strand-exposed states, in particular to the partially A-strand-exposed state; (2) the similarity between the WT intermediate-pressure partially unfolded state (IP-UNF) and the partially unfolded W2AW4A state, suggesting that IP-UNF corresponds to a sparsely populated partially unfolded ambient-pressure WT state which is in slow exchange with the folded fully A-strand-exposed state; and (3) pressure-dependent fast exchange between folded and unfolded WT states, suggesting kinetic accessibility of partially unfolded WT states.

Trp4 exposure in isolation only marginally increases the very fast exchanging random coil population in the BC loop; the combined exposure with Trp2, in contrast, leads to a dramatic increase of both a very slow exchanging partially unfolded state and local very-fast-exchanging RC population. We also conclude that any local destabilization of the BC loop region by Trp4 occurs via the very slow exchanging partially unfolded state. A higher population of the partially unfolded state may also promote formation or stabilization of the folded fully A-strand-exposed state. These newly discovered partially unfolded states and local unfolding, in conjunction with strand-swapping, provide the kinetic framework to stabilize dimer-strand-binding over self-insertion of the strand, as described in other domain-swapping proteins (Byeon et al., 2004, Esposito and Daggett, 2005, Liu et al., 2012, Gronenborn, 2009, Spadaccini et al., 2014, Liu and Huang, 2013, Kang et al., 2012).

Modulation of Cadherin-11 monomer stability could, in principle, be fine-tuned by a dimerization partner or by intermolecular contacts in the X-dimer. Notably, the contact surface in type II Cadherins is particularly large and hydrophobic, compared to type I cadherins. Similar to other strand-swapping proteins, partially unfolded regions or hydrophobic surfaces could support encounter complex formation or be viewed as “protein solvent” to facilitate domain swapping, conceptualized as an intermolecular folding process. At high concentrations, as encountered in the 2D-network of Cadherins organizing to form adherens junctions, encounter complexes involving partially unfolded proteins might be advantageous.

According to the Le Chatelier principle, for a single slow dimerization process, the relative populations of monomer states participating in exchange processes faster than dimerization are not concentration dependent. Therefore, the concentration dependence of the observed chemical exchange processes implies not only that the partially strand exposed state is on pathway for Cadherin-11 dimerization, but also that more than one dimerization pathway exists ***(Fig. S13b)*** (Miloushev et al., 2008).

The present work has revealed a complex hierarchy of sparsely populated conformational states of the Cadherin-11 EC1 domain. These states, ranging from localized alternative conformations of loops surrounding the Trp binding pockets to partially or fully A-strand exposed states (with exposure of Trp2, Trp4 or both) to more extensively partially unfolded conformations, are a central aspect of Cadherin-11, and probably other type II Cadherin, function. Trp4 is present in type 11, but not type I cadherins. We found Trp4 exposure in the fully A-strand-exposed state to give a larger partially unfolded state population and local destabilization at least of the FG loop and G strand region. In addition to the common advantages of unfolded intermediates in domain-swapping proteins, unfolded intermediate states might offer an evolutionary advantage for fine-tuning differential Cadherin dimerization specificity.

## Acknowledgements

We acknowledge Shibani Bhattacharya (New York Structural Biology Center, NYSBC), Mike Goger (NYSBC), Julia Brasch (Columbia University) and Anna Kacyznska (Columbia University) for helpful discussions. This work was supported by National Institutes of Health grant R35 GM130398 (A.G.P.), by National Institutes of Health grant R01 MH11481708 (L.S.) and by the US National Science Foundation grant MCB-1412472 (to B.H.). Some of the NMR experiments were conducted at the Center on Macromolecular Dynamics by NMR Spectroscopy located at NYSBC, supported by a grant from the NIH National Institute of General Medical Sciences (GM118302). B.H., L.S., and A.G.P. are members NYSBC. The data collection at NYSBC was made possible by a grant from ORIP/NIH facility improvement grant CO6RR015495. Data collected using the 800 MHz Avance III NMR spectrometer is supported by NIH grant S10OD016432. The 700 MHz NMR spectrometer was purchased with funds from NIH grant S10OD018509. The 900 MHz NMR spectrometers were purchased with funds from NIH grant P41GM066354 and the New York State Assem bly.

## Author Contributions

B.H., L.S., and A.G.P. conceptualized the research program. H.K. and A.G.P. designed NMR experiments. H.K. conducted research, wrote the data analysis scripts, and analyzed the data. All authors contributed to interpretation of the results and writing of the paper.

## STAR Methods

### RESOURCE AVAILABILITY

#### Lead Contact

Further information and requests for resources and reagents should be directed to and will be fulfilled by the lead contact, Arthur G Palmer.

#### Materials Availability

This study did not generate new unique reagents except plasmids and proteins (see *Protein expression and purification)*. Plasmids are available upon request.

#### Data and code availability

The commented code for multi-site fitting of ^15^N-relaxation dispersion data and relaxation dispersion data were deposited at Github: https://github.com/hanskoss/2021Structure_CadhMultisite.Key results of the fitting process are listed in ***Tables S3-S5***.

The ^1^H, ^15^ N backbone resonances for all constructs were only partially assigned. Important chemical shifts for some constructs under some conditions are listed in ***Tables S1-S2***.

## METHOD DETAILS

### Protein expression and purification

Plasmids for overexpression of WT Cad11-EC1 (residues 1-98) and Cad11-EC1EC2 (Residues 1-207) were constructed using a pSMT3 vector (Patel et al., 2006). The constructs included an N-terminal 6-HIS-SUMO-tag. WT plasmids were generously provided by Julia Brasch and Anna Kaczynska (L.S. lab, Columbia University). Plasmids for W2A, W4A and W2AW4A mutants were obtained commercially (Genewiz).

The following procedure was used to produce ^15^N or ^13^C, ^15^N-labelled protein. The pSMT vector harboring the construct was transformed into E. coli strain C41 (DE3) on 2xYT/Kanamycin plates. Eight colonies were picked and incubated to 37 °C overnight in 5 ml 2xYT/Kanamycin. Glycerol stocks were prepared from the overnight cultures. Cultures were inoculated from glycerol stocks and grown until plateauing at high density (OD∼ 4) in 1-3.5 l 2xYT/Kana (0.5 l/flask) at 37 °C. Following a protocol related to Murray et al. (Murray et al., 2012), cells were pelleted and carefully resuspended in a 2l flask with 500 ml minimal medium (90 mM Na_2_HPO_4_, 22 mM KH_2_PO_4_, 8.5 mM NaCl, 18.7 mM ^15^N-NH_4_Cl, 28 mM D-Glucose, 8.7 Mm Na_2_SO_3_, 0.5 x Trace Metal Mix, 50 µg/ml Kanamycin, 20 mg/l Thiamin, 20 mg/l Biotin, 2 mM MgCl_2_, pH 7.4, 3 mM CaCl2). The glucose concentration was set to 11 mM for ^13^C-labelled proteins (substituting regular glucose with D-Glucose-^13^C _6_). The flask was kept at 37 °C for 45 minutes and then cooled to 20 °C for 30 minutes. Protein expression was induced with 100 µM IPTG. Bacteria were pelleted 10-14 hours after induction, and frozen at -80 °C. For expression in 100% D_2_O, an initial pre-culture from glycerol stock was grown in 5 ml 2xYT/Kanamycin media prepared in D_2_O. The minimal medium for deuterated protein (2H-MM) production was modified by using D_2_O, anhydrous reagents and reagents dissolved in D_2_O (exception: H_2_O in HCl which is used for FeCl_3_ solution preparation); glucose was not deuterated. In a typical procedure, the pre-culture was transferred to 500 ml ^2^H-MM and grown to an OD of ∼1.5. After centrifugation, the pellet was carefully resuspended in 2l ^2^H-M M, distributed between 4 21-flasks. At OD 0.5-0.6, the incubation temperature was lowered from 37 °C to 20 °C, followed by induction (100 µM IPTG) 30 minutes later. Bacteria were pelleted after 18-24 hours and kept frozen -80 °C.

All purification steps were performed at 4 °C. 20 ml lysis buffer (25 mM TrisCI, pH 8.0, 250 mM NaCl, 40 mM imidazole, 10 mM benzamidine, 1 mM MgCl_2_, 3 mM CaCl_2_, lysozyme) and one tablet complete, EDTA-free protease inhibitor tablet (Roche) were added to frozen pellets in large diameter centrifugation vessels (1000 ml culture per vessel). The vessels were placed on a shaker (120 rpm) for 30 minutes. A small amount of powdered DNAse l (bovine pancrease) was then added before shaking for another 30 min and centrifuging for 30 minutes (3210 g) to remove the bulk of larger cellular remnants. The supernatant was used for further purification using the AKTA Purifier system (GE Healthcare). A HisTrap crude (GE Healthcare) column and His buffers A (25 mM TrisCI, pH 8.0, 500 mM NaCl, 40 mM imidazole, 1 mM TCEP, 3 mM CaCl_2_) and B (25 mM TrisCI, pH=8.0, 500 mM NaCl, 500 mM imidazole, 1 mM TCEP, 3 mM CaCl_2_) were used for the first purification step. The column was washed with at least 300 ml His A prior to elution.

The SUMO tag was cleaved by adding 100 - 500 µl of 1 mg/ml ULP1 protease for at least 20 hours, with the ULP1 concentration adjusted to achieve near complete cleavage. This was followed by at least two 3-5 hour dialyses (MW 10 kDa tubing) steps against His Chelating Buffer (25 mM TrisCI, pH=8.0, 250 mM NaCl, 20 mM imidazole, 1 mM TCEP, 3 mM CaCl_2_) with 1 mM PMSF (stock : 0.2M in isopropanol). For the second purification step, the sample was run over a HisTrap Chelating column (GE Healthcare) to collect the tag-free flow-through. The collected sample was dialyzed (MW 10 kDa tubing) against Low Salt buffer 2x for at least 3-5 hours (25 mM TrisCI, pH 8.0, 1 mM TCEP, 3 mM CaCl_2_). Size exclusion chromatography used a Hiload 16/60 Superdex 76 PG (Fisher Scientific) column. The sample buffer used for gel filtration is composed of a pressure resistant mixture (Quinlan and Reinhart, 2005) of 10 mM TRIS/tricarballylate (pH = 7, 6.905 Tris, 3.095 mM tricarballylate), 150 mM NaCl, 10% D_2_O, 3 mM CaCl_2_, followed by spin concentration using Amicon Ultra 15 Centrifugal Filters (10k kDa). Final samples contained 0.02 % (w/v) NaN_3_ and a protease inhibitor mix (stock: 100x in 10% DMSO, end concentrations 0.5 mM AEBSF, 10 µM E-64, 2 mM Benzamidine; a generous gift from Eric Greene, Columbia University).

### NMR data acquisition

#### general aspects

NMR data were acquired with Bruker 500-, 600-, 700-, 800- and 900-MHz spectrometers with triple-resonance z-gradient cryogenic probes (600 MHz at Columbia University, all others at New York Structural Biology Center). DSS either in the sample or separately in sample buffer was used to confirm correct temperature calibration.

#### 2D spectra

For most TROSY-selected experiments in this work, an experimental TROSY-selected experimental framework corresponding to lgumenova et al. was used (lgumenova and Palmer, 2006). In order to enable a short recycle delay of 800 µs, the experiment was performed in a BEST-manner with an PC9,4,90 (first π /2), Eburp2, time-reversed Eburp2 π /2, and Rsnob π pulses (Lescop et al., 2010). 2D ^15^N-HSQC experiments were recorded either as TROSY-selected, BEST-type TROSY, regular TROSY or SOFAST experiments. Note that the 2D spectrum in Fig. 2a is the projection sum of a pseudo-3D *R*_*ex*_ (see below) experiment, which accentuates random coil peaks due to prolonged decay of peaks belonging to slower tumbling folded regions.

### 3D backbone resonance assignment experiments

For backbone resonance assignment of C11-WT, TROSY versions of the HNCA, HNCOCA, HNCACB, HNCACO, HNCO and HNCACO were used as 3D experiments. For C11-W2AW4A, only HNCA, HNCO, and HNCACB experiments were recorded, for C11-W4A, only HNCA was recorded. All determined ^1^H, ^15^N - chemical shifts are listed in ***Table S1***.

#### ^15^N-R_2_/R_1_ ratios

To record ^15^N-*R*_*2,0*_/*R*_*1*_ ratios, ^15^N-*R*_*1*_ and ^15^N-*R*_*2,0*_ (exchange-free) were recorded for an 80 µM sample at 285 K and 298 K. ^15^N-_*R1*_ was recorded as a TROSY experiment using flip-back pulses and a Watergate scheme (Zhu et al., 2000). Relaxation delays were 24 ms, 336 ms (2x for error determination), 656 ms, and 976 ms. Data were recorded in an interleaved manner, 180 points in ^15^N dimension, total acquisition times: 1498 min (285 K), 967 min (298 K).

For exchange-free ^15^N-*R*_*2,0*_, the experimental TROSY-selected framework (see above “2D spectra”) was modified to include a relaxation period T. To suppress chemical exchange, a spin-lock was applied during T with a central S3E element dividing the two T/2 block. The experiment was set up in an interleaved manner, recording for each of the 200 ^15^N points the following experiments: 4 x [3 x [T=15 ms with spin lock (power: 1 kHz); T=15 ms us with no spin lock]; 1 x [T=15 ms with spin lock (power: 1 kHz); T = 0 us]]. The 16 experiments with spin locks had the following offsets: 2 × 104.5 ppm; 2 × 108.5 ppm; 2 × 112.5 ppm; 2 × 116.5 ppm; 2 × 120.5 ppm; 2 × 124.5 ppm; 2 × 128.5 ppm; 2 × 132.5 ppm. To minimize off-resonance artifacts, any peaks shifted > 4 ppm (^15^N) from the spin lock center was discarded. Therefore, per ^15^N point, 12 spin-lock-free relaxation experiments for T=15 ms were obtained; 4 experiments with T=0 µs, and 4-6 spin-lock experiments with T=15 ms for any given residue. To obtain *R*_*2,0*_, we calculate:

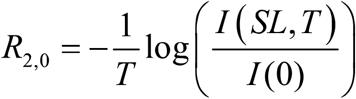

In this case, the exponential decay between T=0s and T=15 ms with spinlock had to be obtained; data with T=15 ms without spinlock were not needed. Note that the spinlock with B_1_ = 1 kHz was not sufficient to suppress all exchange, probably accounting for the elevated *R*_*2,0*_ values around residue Asp32.

#### ^15^ N-Rex experiments

Experiments to obtain ^15^ N-*R*_*ex*_ were executed at 285 K using the scheme described for obtaining exchange-free ^15^ N-*R*_*2,0*_ with varying offsets and spin lock power levels. As a representative example, for each ^15^ N point interleaved experiments were recorded in the following order: SL, SL, noSL, SL, SL, noSL, SL, SL, noSL, SL, SL, noSL, SL, SL, TO, TO (SL: with spin-lock; noSL: without spinlock; TO: TROSY), with SL offsets spread out over the ^15^ N spectral width. *R*_*ex*_ was obtained from

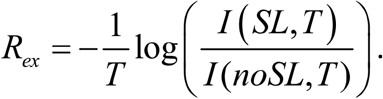

The experimental conditions for the data sets recorded for global fitting are listed in ***Table S6***.

Error estimation: The average of the square root of the squared standard error of the absolute intensity for each residue was obtained. This averaged error was then propagated to *R*_*ex*_ individually for each residue using Monte Carlo simulations. We found that residue-specific differences in error mostly stem from absolute intensity differences, and those are preserved in our approach.

#### ^15^ N-CPMG experiments

The TROSY-selected framework used for the ^*15*^ N-*R*_*2,0*_ and *R*_*ex*_ experiments was modified by replacing the spin-lock block with constant-time CPMG blocks. Each half of the CPMG block, divided by an S3E-element, contains N repetitions of 4 π pulses (phases: x, x, y, -y), separated by variable τ _cp_ For the various combinations of N and τ _cp_, constant-time (total time T) CPMG blocks were recorded in an interleaved manner, with the objective to average heating by not sequentially recording two experiments with similar high (short τ _cp_) or low (long τ _cp_) heating.

The experimental conditions for the data sets recorded for global fitting are listed in ***Table S7***.

In analogy to our error estimation for *R*_*ex*_ method but having the benefit of multiple data points per curve, we obtained average absolute intensity errors for each curve collected from all τ _cp_ values for which more than one τ_cp_ point had been recorded. The error for each intensity ratio (I_T_/I_0_) and subsequently R_cpmg_ was then calculated using standard error propagation. The positive and negative errors were calculated separately unless the negative error calculation led to negative intensity ratios.

#### ^15^N-CEST experiments

Some ^15^N-CEST experiments were based on the Bruker library pulse sequence hsqc_cest_etf3gpsitc3d (15/09/22), which is a non-TROSY sequence with decoupling during acquisition. Alternatively, a TROSY- selected version of the experiment was run, based on the framework for ^15^N-*R*_*2,0*_ and *R*_*ex*_. During the CEST period T, CPD decoupling was employed in the proton channel. All experiments, including various CEST offsets and reference experiments were run in an interleaved manner at 285K.

For the 160 µM sample, three non-TROSY experiments at 700 MHz with T = 400 ms for three different B1 fields, 15 Hz, 25 Hz, and 50 Hz, were recorded. For the 640 **µM** sample, the TROSY-experiment was used with T = 400 ms and B_1_ = 35 Hz.

For further processing, ^15^N-CEST data were trimmed to reduce the number of information-free data points. Separately, sections with frequency distant from the CEST dip were used to define reference CEST intensities and to obtain the intensity errors. CEST data points< 0.5 ppm away from the CEST dip minimum were discarded for the CEST analysis.

#### ^1^H-CPMG experiments

^1^H-CPMG experiments were performed described following the pulse sequence provided by Li et al. (Li et al., 2013). Due to a relatively fast degradation an of the deuterated sample which was used for the experiments, the total acquisition times per sample were of limited duration, giving large error bars. Uncertainties in the concentrations an imperfect comparability of temperature between experiment limit the recorded data to be used for qualitative purposes only. For aspects of initial error calculation, see the section about ^15^N-CPMG experiments.

#### High pressure experiments

The Daedalus Xtreme-60 apparatus was used for static high pressure NMR experiments. Mineral oil was used as the buffer fluid for the pressure tubes. The standard air removal procedure was repeated three times prior to connecting the sample. For titration experiments, the experiments were started at a high pressure and then reduced stepwise. To rule out problems with equilibration, control experiments, either by repeating by changing the direction of pressure change, were performed. We found that in our NMR buffer system, pressure hardly affects the ^1^H reference shift.

### NMR data processing

#### General approach

Raw NMR data were pre-processed using customized bash scripts and NMRPipe (Delaglio et al., 1995). CCPN analysis 2.5.0 (Vranken et al., 2005) was used for peak assignment and extraction of peak intensities and chemical shifts. For peaks with particularly small intensities, e.g. at noise level, the peak fitting algorithm tends to overestimate peak intensities. For that reasons we decided to obtain peak intensities by adding the sum of all spectra from any pseudo-3D dataset to each individual spectrum for fitting at the preprocessing stage with NMR Pipe. The peak intensities obtained from the sum of all spectra are then subtracted from the sum of all spectra + spectrum of interest after peak fitting with CCPN analysis. Depending on the experiment type, specific data formats were established, also to include all relevant experimental conditions, to ensure smooth automated post processing; conversion bash scripts were used to avoid any errors and reveal/fix any inconsistencies in the peak lists obtained from CCPN Analyis. Python was used for any subsequent data processing. Errors for data points were calculated as described in the NMR data acquisition section.

#### multi-site fitting process

##### Python scripts for global fitting

We used Python scripts for data processing (https://github.com/hanskoss/2021Structure_CadhMultisite). Documentation provided with the scripts outlines the data model used to combine data from ^15^N-CPMG, ^15^N-CEST and ^15^ N-R_ex_ experiments, at various fields (500, 700, 800 and 900 MHz) and concentrations (160 µM and 640 µM), for a single global fit.

##### Preliminary tests and establishment of boundary conditions

Initial tests with wide boundary conditions and inspection of the relaxation dispersion datasets established the requirement to use at least a three-site model, including a fast and a slow exchange process. We found that the appropriate scheme fast exchange between sites A and B and slow exchange between sites A and C, with A being the major site. To limit the parameter number in the global fit, we assumed *R*_*2,0*_ and Δ*ω*_*AB*_ and Δ*ω*_*AC*_ to be identical for different concentrations. The initial fit gave broad result ranges for parameters, specifically for all Δ*ω* values and *k*_*ex*_ which are known to be highly correlated. We tested a variety of Δ*ω* restraints. We also found that the Δ*ω* of many residues were highly correlated for Δ*ω*≫ 0 (e.g. Δ*ω*_*AC*_ of residues 38 and 14). Therefore, fixing any of the Δ*ω*≫ 0, of which there are a total of 2*(number of residues), greatly shrinks the parameter space offitting results. Initially, we attempted to set Δ*ω*_*AC*_ to known chemical shift perturbations: The chemical shift position for the main state A was set to be the position of the recorded peak which is justified by the fact that the corresponding exchange process is generally in the slow exchange regime. Chemical shift for position C was set in the following ways: (a) chemical shift of other states or mutants, specifically the unfolded state, the folded state of the W2A mutant, the folded state of the W4A mutant, the folded state of the W2AW4A mutant, and the dimer state; (b) The sign of the chemical shift was assigned from CEST curve if a clear asymmetry was visible. For W2A, there were no conflicts between (a) and (b). The following residues were therefore restrained based on W2A shifts: 30, 32, 43, 45, 53, 54, 55, 73, 77, 78, 86. For residues 38,73,77 and 86, the sign of the chemical shift was inferred directly from the shape of the CEST curve. Δ*ω*_*AC*_ for residues 37 and 50 was set freely.

For residues Gly43 and Gly55, very large, unrealistic *Δ*_*B*_ emerge under any scenario leading to reasonable *χ*^*2*^ for these residues. Inspection of the CPMG curves and *R*_*ex*_ data suggests partial quenching of a separate very fast exchange process which is either unique to these residues or equivalent to one of the other processes discussed in the paper, possibly originating from the fact that the Gly43 and Gly55 peaks are in the random coil region. We therefore excluded Gly43 and Gly55 from further analysis.

The following chemical shifts as were established as restraints for the main calculation: site A: WT-Cadll; site B: folded W2AW4A-Cad11 and unfolded W2AW4A-Cad11 shifts as boundaries, but only for loop BC, DE and FG residues; site C: W2A-Cad11. Exchange between minor sites B and C is possible but hardly changes *χ*^*2*^. Examples for other non-fitting restraints which do return higher *χ*^*2*^ include setting the following chemical shifts: site C to W4A-Cad11, site C to WT-dimer, site B to W2A-Cad11 or site B (generally or for the specified regions) to unf-W2AW4A-Cad11. Setting site B to fold-W2AW4A-Cad11 for loopl-3 increases *χ*^*2*^ moderately.

##### Global fitting procedure

For fitting of ^15^N-CEST data, we transformed the corresponding exact equations for triangular exchange in R_1ρ_ experiments (Palmer and Koss, 2019); for initial calculations, *R*_1ρ_ multi-site approximations were used (Koss et al., 2017). Data from the flat sections of plateau are used for normalization and discarded for the actual fit. A 0.5 ppm region around the main state is removed due to low information content and expected inaccuracies of the model for that region. To calculate the expected position of the CEST dip, the position for all states being in fast exchange with the main state are combined (population-weighted). For fitting CPMG data in stages 1 and 2, the approximations for **H**_22_ from Eq. 18 and for λ_2_ from Eq. 20 in Koss, Rance and Palmer were used (Koss et al., 2018). For stage 3 and 4, the exact equation is used. Theoretical *R*_*ex*_ are by calculating the low-power relaxation rate from the corresponding exact CPMG equation, and then subtracting the high-power relaxation rate calculated from the exact *R*_1ρ_ equation. *R*_*2,0*_ is not a required parameter for calculating theoretical *R*_*ex*_ data. *R*_*2,0*_ is assumed to be equal or larger with increasing **B**_**0**_, which is a useful restriction for fitting CPMG data. *R*_*2,0*_ boundaries in ^15^N-CEST experiments only have a negligible influence on the results and were used to help initial convergence. R_1_ also has only a minor effect in CEST calculations and was set to be constant for the entire protein to reduce the number offitted parameters. We estimated an average *R*_1_ for 700 MHz and 800 MHz from calculations with Hydro-NMR (Garcia de la Torre et al., 2000).

Global fitting is performed in four stages. In stage 1 and 2, starting parameters are randomized within boundaries: for stage 1, those boundaries are derived from the preliminary fitting tests (above); for stage 2, the boundaries are derived from stage 1 results. We observe that Δ*ω*_*B*_ and Δ*ω*_*C*_ generally cannot cross 0 during fits. For stage 2, the sign of either 10 or (with reasonable rest rictio ns, *vide infra)* 4 sets of Δ*ω*_*B*_ and Δ*ω*_*C*_ were therefore randomized (1024 or 16 possibilities, respectively). In stage 2a, 300 fits with 9 steps (documenting progress every 3 steps) were initially performed. The 100 fits with the lowest *χ*^*2*^ were then continued (stage 2b), with 50 steps (documenting progress every 5 steps). Stage 2b results were used to explore convergence of the global fit and judge the number of parameter set clusters with small *χ*^*2*^. Elsewhere in the paper we conclude that the field-dependent chemical shifts are dominated by the process between the major site and state B; this is supported by the observation that the boundaries for Δ*ω*_*B*_ of residues 30, 32, 54 and 78 align with the observed field-dependent chemical shift. We therefore further restrict the signs for Δ*ω*_*B*_ by considering field-dependent chemical shift direction and, for Gly45, the fact that a positive Δ*ω*_*B*_ would give an unlikely large δ_N_ We repeated stages 2a and 2b with the additional restrictions and selected the smallest *χ*^*2*^ result (all results were very similar), and then randomized the sign of the four ambiguous Δ*ω*_*B*_ and Δ*ω*_*C*_, giving a total of 16 starting sets, which we then minimized (50 steps, stage 3). The average of the resulting parameters are the main results. For error calculation (stage 4) we used all 16 resulting parameter sets.

##### Error estimation and concentration differences

After obtaining the global fit solution, the error was estimated using bootstrapping (stage 4), with 5 calculations for each of the 16 results from stage 3 ***(Tables S3-S5)***. For that purpose, for each of the data sets (individual CEST, CPMG curves, all *R*_*ex*_ per residue), residuals *δ*_i_ = (y_i,fit_ - Y_i,exp_)/y_i,err_ were calculated. The lists of residuals were drawn randomly with replacement to create new datasets with *δ*_j,new_ = y_j,fit_ + Y_j,err_ * *δ*_i_. Normalization by the errors was used to account for heteroscedasticity. The new datasets were then globally fit (50 steps) to obtain alternative sets of parameters (main results as starting parameters). Multiple repetitions of this process were used to produce parameter ranges of la uncertainty. For each of the 80 calculations, the ratio and the difference between key parameters *P*_b_, *P*_c_, *k*_12_+*k*_21_ and *k*_13_+*k*_31_ at high and low concentrations was calculated to test whether a general concentration dependency of these parameters emerges. In an alternative calculation, we permitted exchange between the sparsely populated sites ***(Tables S3-S5***, ***Fig. S13a)***.

##### Corroborating Δω restraints

We tested the *Δω* restraints with all BC, DE and FG residues, including data with almost flat relaxation profiles or with large uncertainties (***Fig. S10***): We used all known W2A and W2AW4A chemical shifts for setting Δ*ω* restraints and compared our obtained model (with fixed *k*_12_+*k*_21_, *k*_13_+*k*_31_, *p*_b_, *p*_c_ and Δ*ω* from known W2A and W2AW4A chemical shifts) to unrestrained runs for each residue. Leaving only Δ*ω* unrestrained did not improve *χ*^2^. Leaving *k*_12_+*k*_21_, *k*_13_+*k*_31_, *p*_b_ and *p*_c_ free for each residue gives only very slightly lower *χ*^2^ than restraining it globally. Allowing chemical exchange between the minor sites B and C does not change *χ*^2^. The model is supported by qualitative analysis of ^1^H-CPMG relaxation data recorded for [U-^2^H, U-^15^ N]-labeled samples (***Fig. S11***).

#### mapping chemical exchange to the structure by evaluating spectra recorded at different B_0_ and temperatures

The methods shown here aims to include relaxation processes encoded in the ^15^ N and ^1^H dimension and peaks with intensities too low for RD analysis. We estimated the degree of slow-intermediate exchange between the main state and the W2A-like (partially strand exposed) state directly from combining knowledge from the RD analysis with peak intensities recorded at different conditions. Fast exchange displays a pronounced temperature and field dependency of peak intensities. Fast exchange at 298K is essentially invisible in RD analysis. For peaks with slow-intermediate exchange, we also find a different field dependency of peak intensities. We therefore calculated the intensity ratio

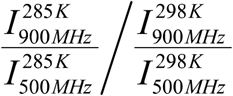

from ^1^H, ^15^ N-TROSY-HSQC spectra and plotted it on the structure. A large ratio maps to a large Δ*ω*_AC_, which corresponds to a site with major exchange between a major A-strand-bound and a sparsely populated partially A-strand-exposed (W2A-like) state.

#### obtaining random coil populations of very fast exchanging residues

The relative population of random coil is, by definition, 100% in the RC state. While it cannot be ruled out that the fraction of remnant random-coil state in the WT-dimer state is ≫ 0%, the fact that its main peak position is well in the folded region of the spectrum suggests that this fraction will not be large. For purposes of comparing relative folded/RC populations for WT-monomer and mutant states, we assume WT-dimer to be the reference state. We observe that, while the various mutants generally follow the order mentioned above, the degree of disorder varies from residue to residue.

#### dependency of chemical shifts and intensities on high pressure

The pressure dependent measurements included intensities and chemical shifts (^1^H, ^15^N) for multiple residues, two states (folded/unfolded) and two temperatures (285K, 298K). The plateau values for the intensities of the most unfolded state were not reached at the highest pressure. For that reason, we extracted the characteristics of the curves using spline functions, fitting intensities and chemical shifts at once, but individually for each residue. The first-order two-point spline function did not give satisfactory fits in many cases. We found that either second-order two-point or first-order three-point spline functions generally resulted in good fits, except for the folded peaks for some residues in the FG loop and for residue Ser54 in the DE loop. For the main analysis we refer to the first order spline fit with three points (two straight lines) which returns a critical pressure for each fit. Intensities were scaled to 1 bar (folded peaks) or 1.72 kbar (unfolded peaks).

#### Fitting intensity profiles to high-pressure WT RC peak intensity profiles

The pressure profiles were smoothed using a Savitzky-Golay filter with window length 7, polynomial order of 1 and no extension. For fitting (scaling) any intensity profile set (RC peaks) to a corresponding WT RC peak profile, both profiles were first normalized. The error function to be minimized by fitting was

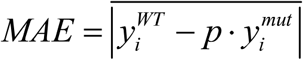

with intensities y for each residue i of the WT or the mutant (mut) construct, optimizing a parameter p. Because the intensities are normalized, 0 ≤ MAE ≤1. With MAE = 0, 100% (1-MAE) of the WT intensity is “explained” by the mutant intensity. Random intensity profiles were generated using a Gaussian normal distribution centered at O and with a standard deviation (scatter) equivalent to the standard deviation for the W2AW4A mutant profile. The fit to obtain an MAE range for random intensity profiles was repeated 500 times, and the MAE interval was then obtained for the desired confidence (sigma) level.

## Supplemental infromation – Supplemental Figures and Tables

**Fig. S1:**
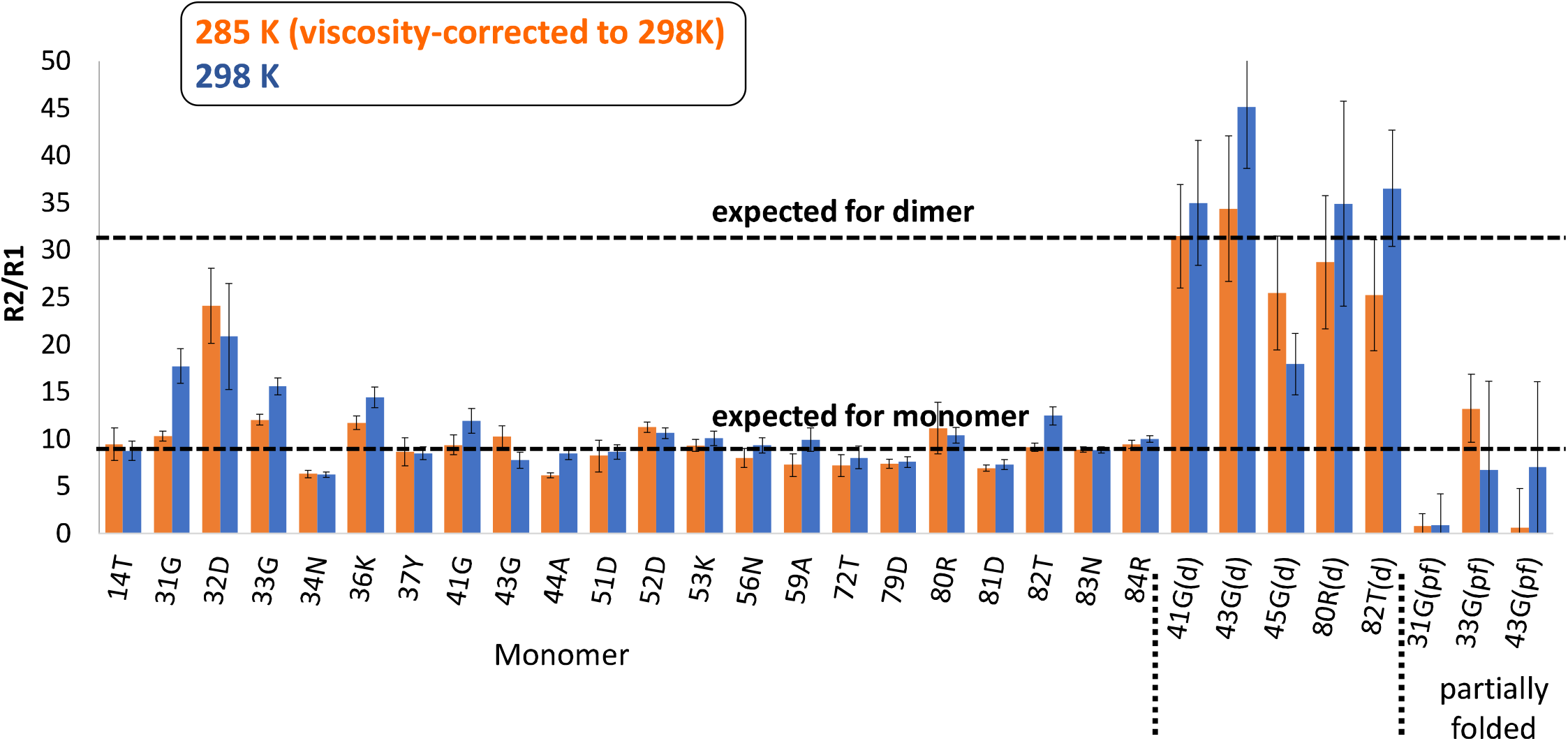
*R*_*2,0*_*R*_*1*_ ratios for WT-Cadherin to establish monomer and dimer status of various peaks. Exchange was suppressed (incompletely for example around residue 32) using a 1 kHz spinlock, and R_1_ was measured by standard methods. *R*_*2,0*_/*R*_*1*_ ratios measured at 285 K were scaled to eliminate the effect of viscosity for comparability with results obtained at 298 K. Theoretical values and the scaling temperature factor were generated with Hydro-NMR.

**Fig. S2:**
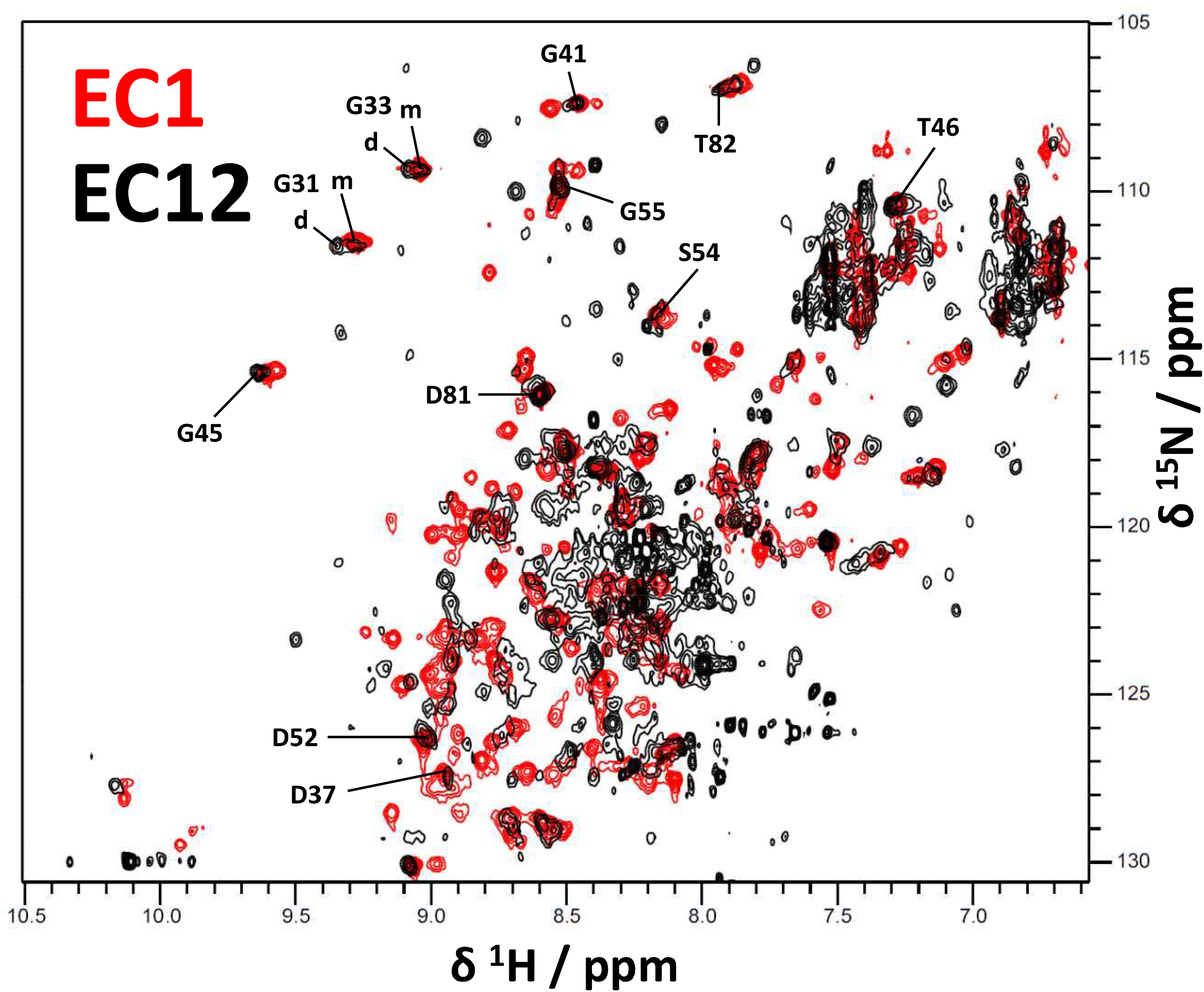
^1^H,^15^ N-SOFAST-HMQC spectra of Cadherin-11-EC1 and Cadherin-11-EC12 domains. Key monomer peaks display a virtually perfect overlap between Cadherin-11-EC1 and Cadherin-11-EC12 domains. Due to degradation and extensive peak broadening, the state populations, and changes in dynamics in the Cadherin-11-EC12 spectrum were not analyzed.

**Fig. S3:**
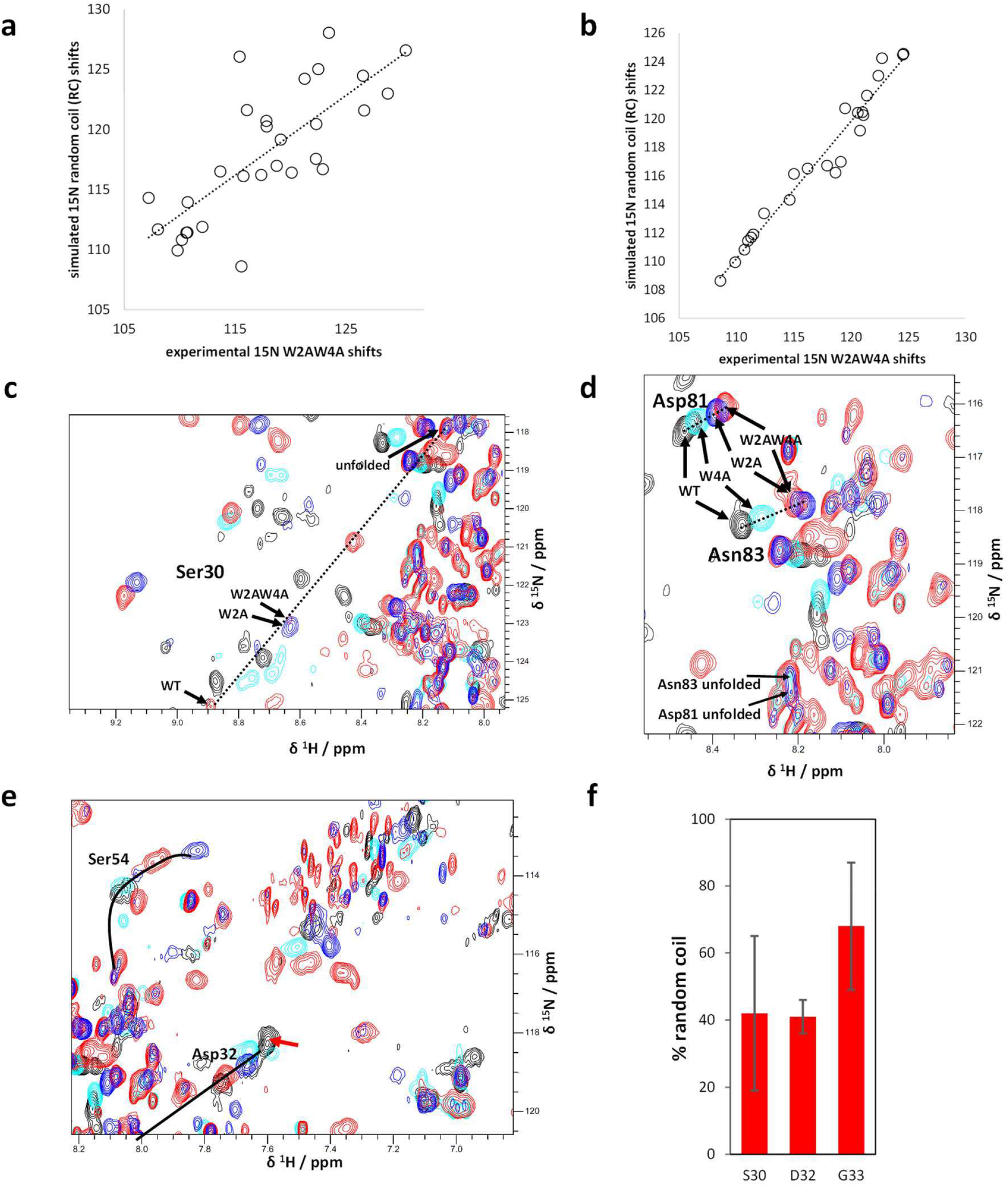
Chemical exchange in Trp2, Trp4 mutants (expanding Fig. 1). **a-c**. Correlation of the “folded” (panel **a**, R^2^ = 0.58) and “unfolded” (panel **b**, R^2^=0.96) W2AW4A-^15^N chemical shifts with simulated random coil shifts confirms that the “unfolded” peaks are random coil (RC). **c-e:** ^1^H, ^15^N-HSQC spectra and peak positions on the fast exchange vector. **f:** W2AW4A-folded RC populations estimated from C_α_ positions. The populations agree with the RC populations estimated from ^1^H, ^15^N spectra (Fig. lc).

**Fig. S4:**
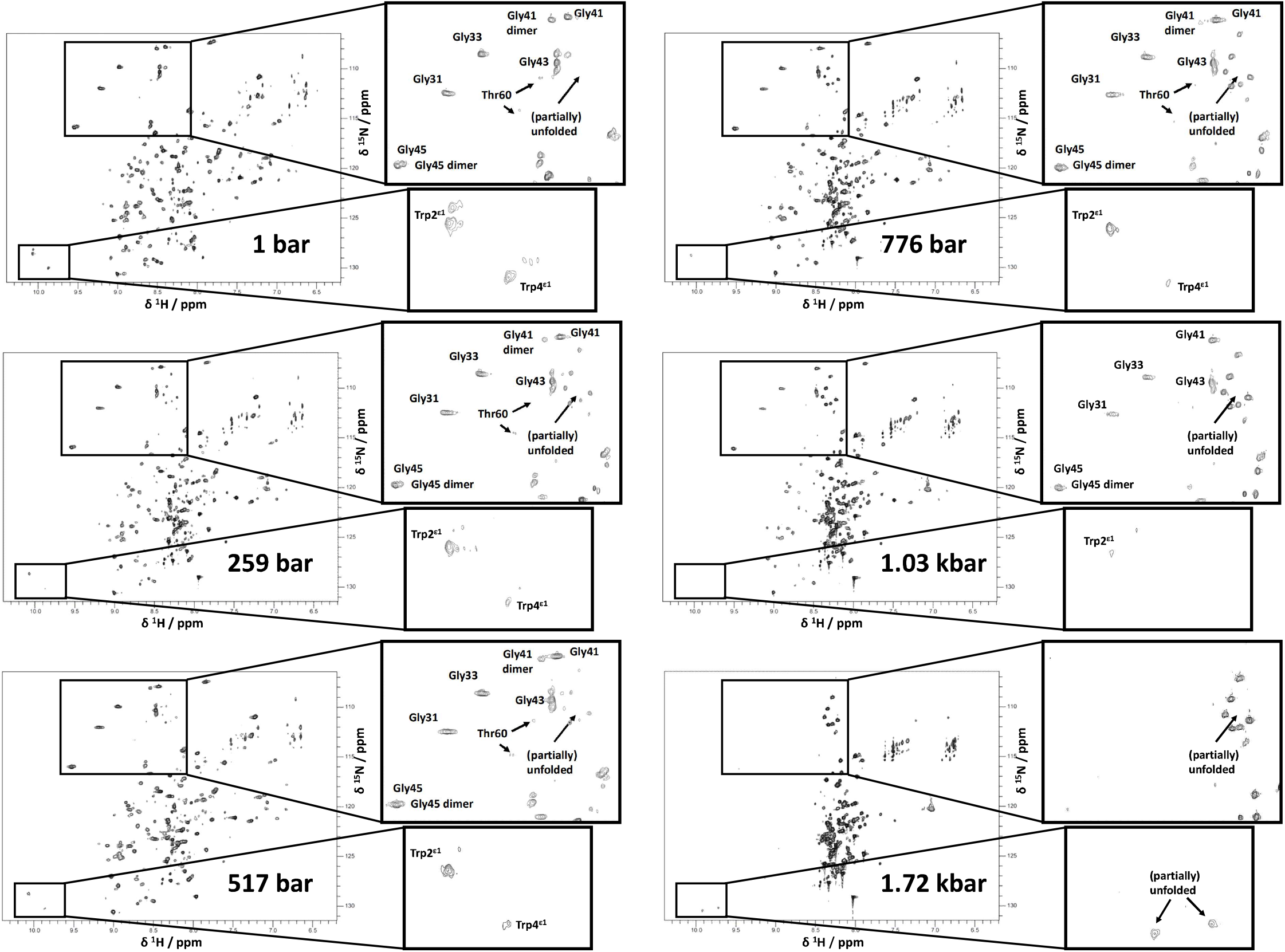
^1^H,^15^N-HSQC-TROSY spectra of Cad11-EC1-WT at different pressures. This pressure series was recorded at 800 MHz, 160 µM and 285 K, the pressures are shown in the figure. The glycine and the Trp side chain regions are expanded using insets.

**Fig. S5:**
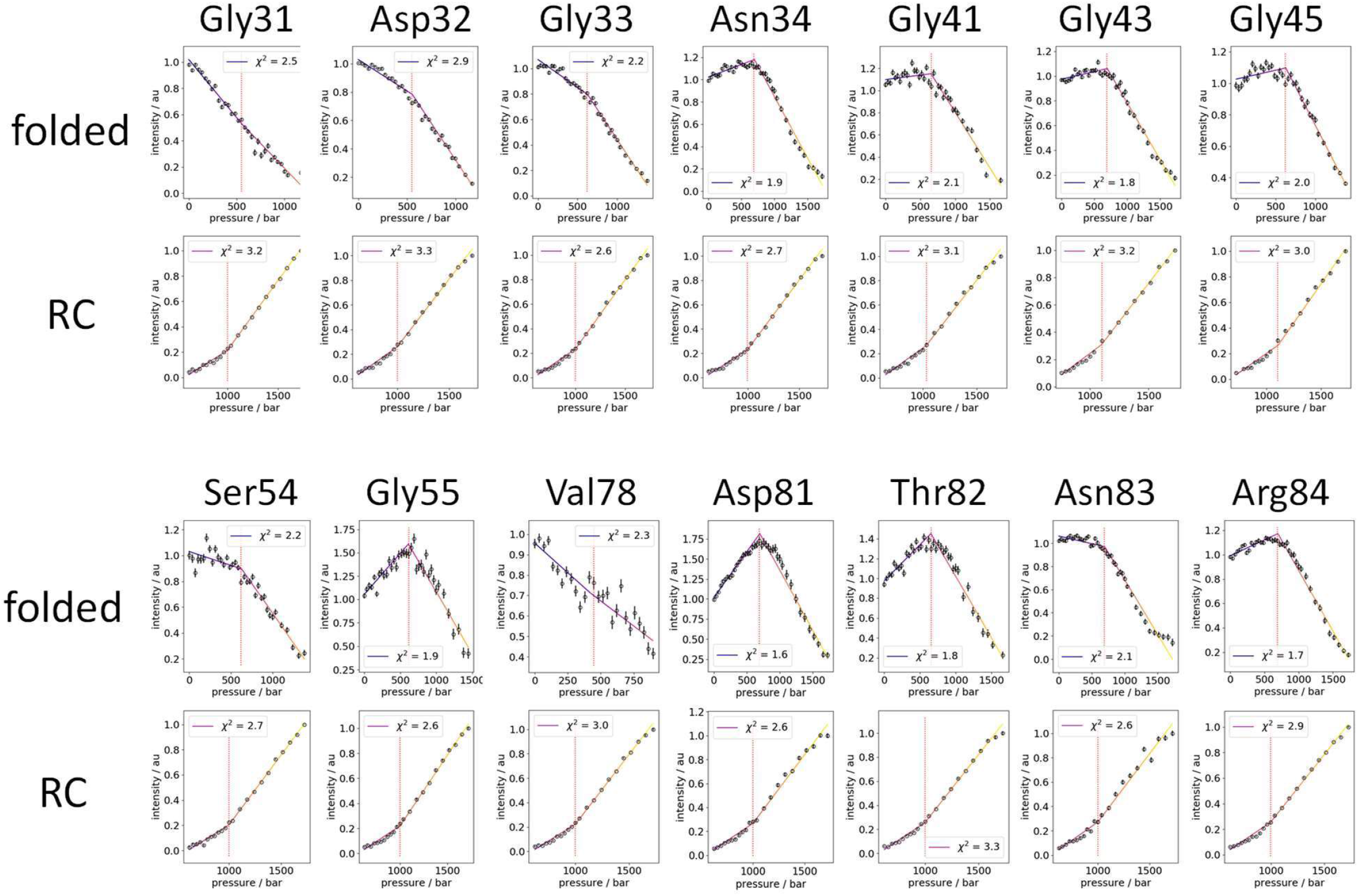
Analysis of pressure-dependency of various peak intensities and shifts at 285K (data and fits). For each residue and the folded and random coil state indicated in the figure, δH and δN profiles are shown. A first order spline with three points (two straight lines) was used to characterize each of the profiles, the results are shown in Fig. S7. The dotted red line indicates the pressure of the central critical pressure for the three-point first order spline fit. χ^2^ is shown in the figure legends. This figure expands Fig. 4a, 4b and 4f.

**Fig. S6:**
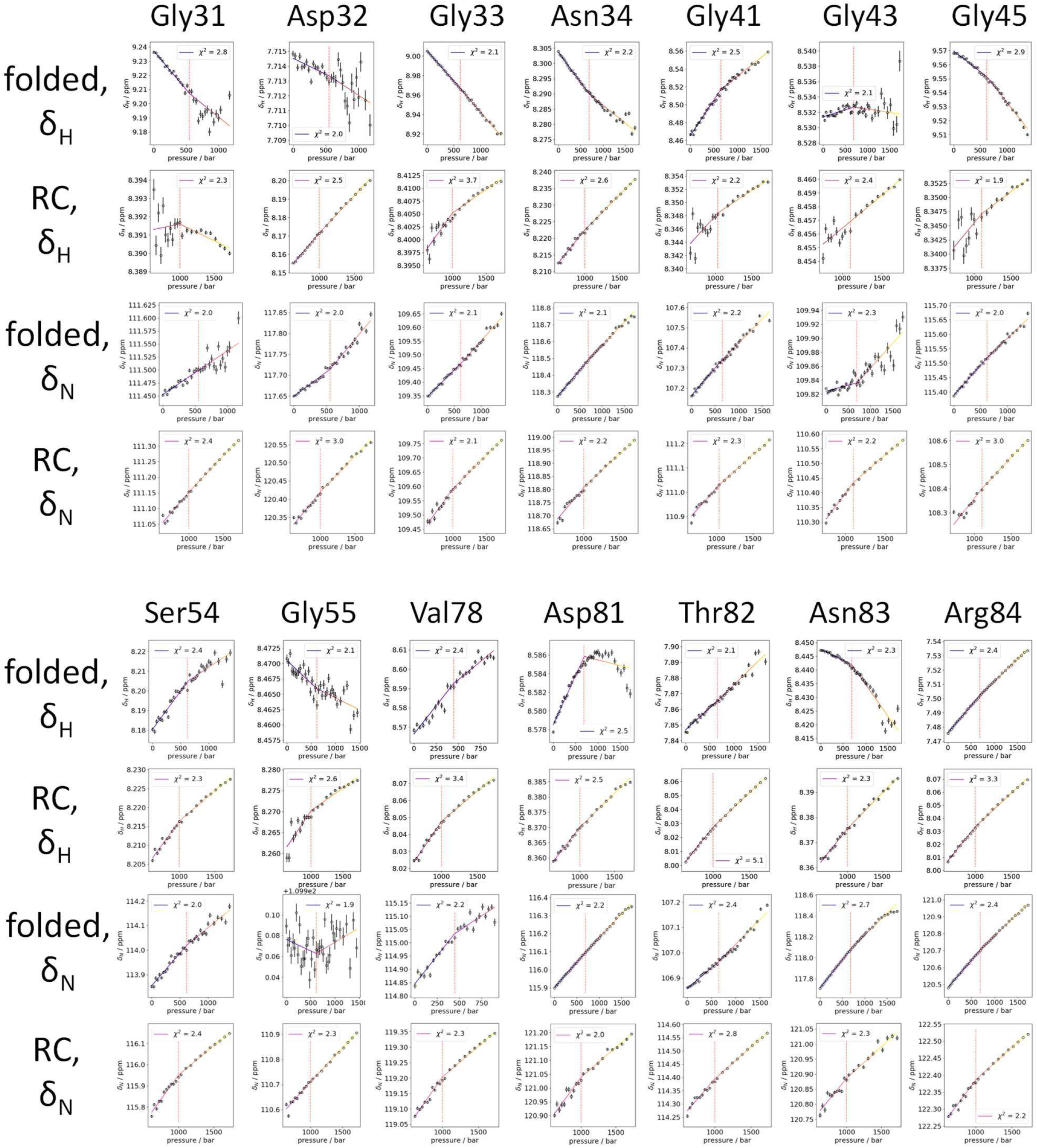
Analysis of pressure-dependency of various peak chemical shifts at 285K (data and fits). For each residue and the folded and random coil state indicated in the figure, δH and δN profiles are shown. A first order spline with three points (two straight lines) was used to characterize each of the profiles, the results are shown in Fig. S7. The dotted red line indicates the pressure of the central critical pressure for the three-point first order spline fit. χ^2^ is shown in the figure legends. This figure expands Fig. 4.

**Fig S7:**
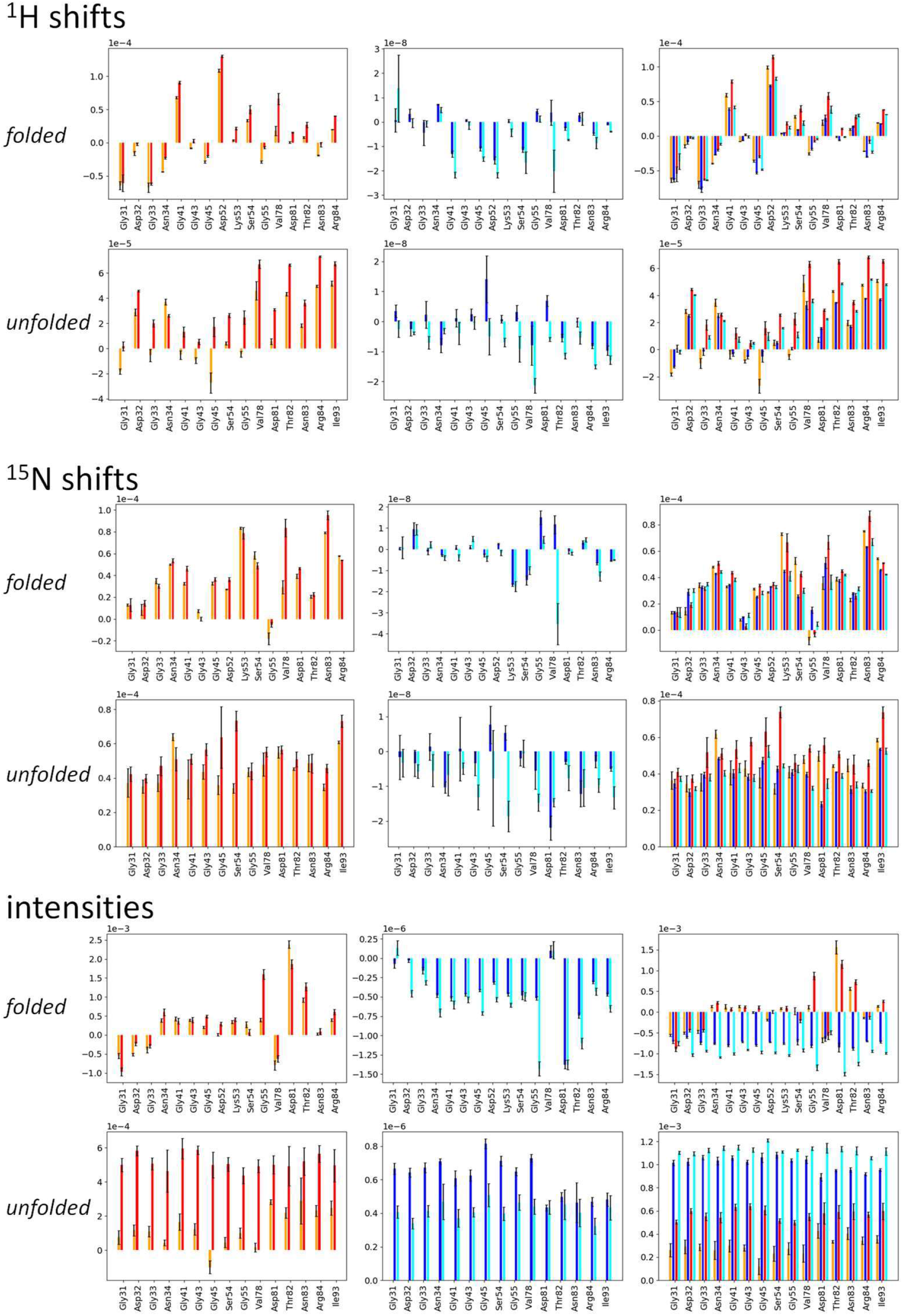
Analysis of Pressure-dependency of various peak intensities and shifts (fitted parameters). Left and center panels: Fitting results for second order two-point spline fits. The slopes at 283K (red) and 298K (orange) are shown in the left panel. The curvatures at 283K (cyan) and 298K (blue) are shown in the center panel. Right panels: Three-point first order spline (right panel, see also Suppl. Figs 5-6) fits give two separate slopes: The low pressure slope for 283K (red) and 298K (orange), and the high pressure slope for 283K (cyan) and 298K (blue). All fitting results are shown for δ_H_ δ_N_ and intensity pressure profiles. Fitting ranges: for folded peaks 0-1.72 kbar, for unfolded peaks 0.62-1.72 kbar. This figure expands Fig. 4c.

**Fig. S8:**
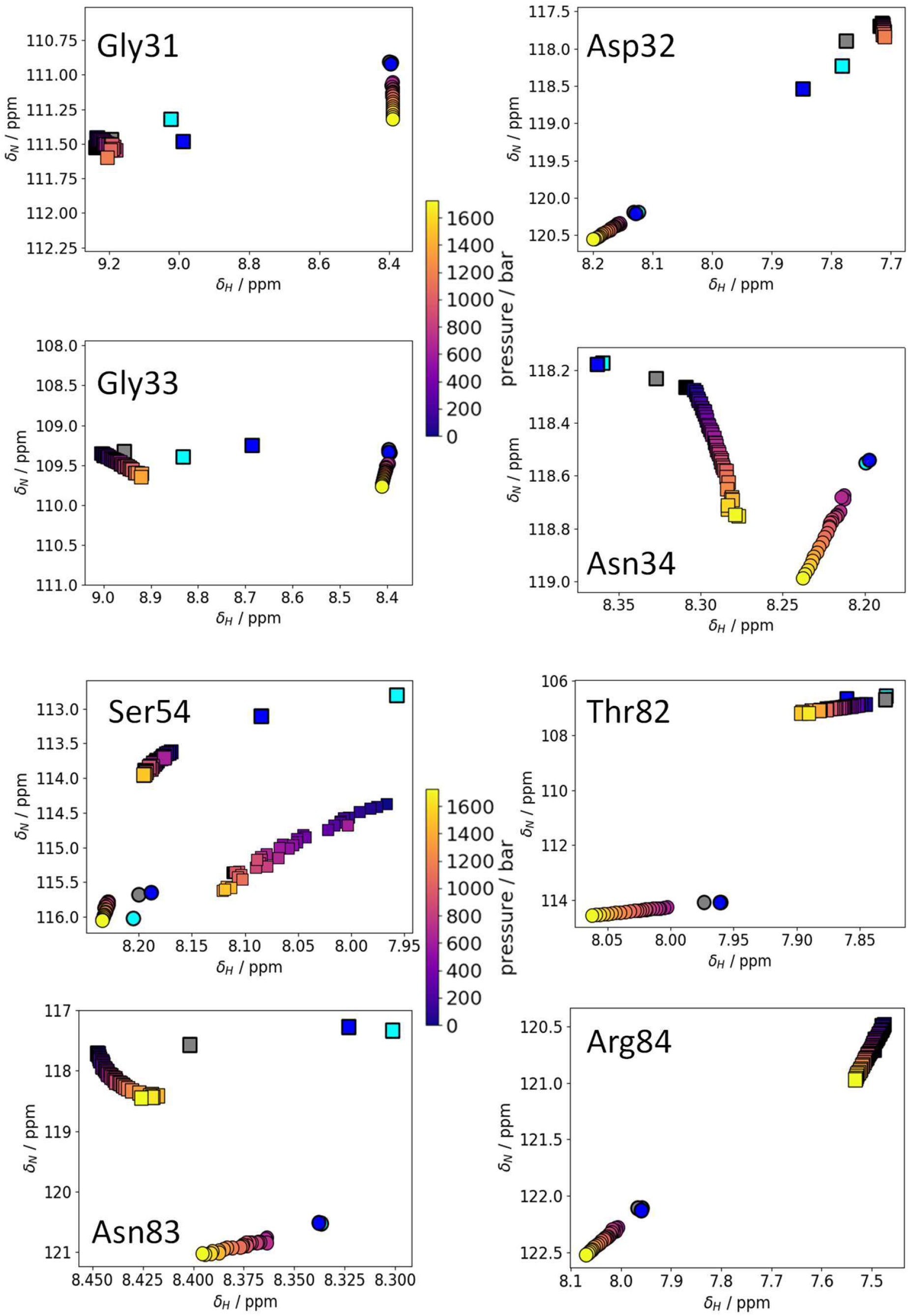
Analysis of pressure-dependent ^1^H, ^15^N-HSQC spectra. For residues in the BC and FG loops, pressure-dependent ^1^H, ^15^N-HSQC spectra are shown expanding Fig. 4g. The color of the calculated curve indicates pressure; the color bar translating colors into pressures is shown. Other symbols: squares - folded peaks; circles - unfolded (RC) peaks; blue: W2AW4A mutant peak; cyan: W2A mutant peak; grey: W4A mutant peak: black: WT peak for comparison with mutants. All data were recorded at 285K, except the high-pressure spectra for Ser54, which was recorded 298K, with the resulting peak pattern displaying a higher curvature than at 285 K.

**Fig. S9:**
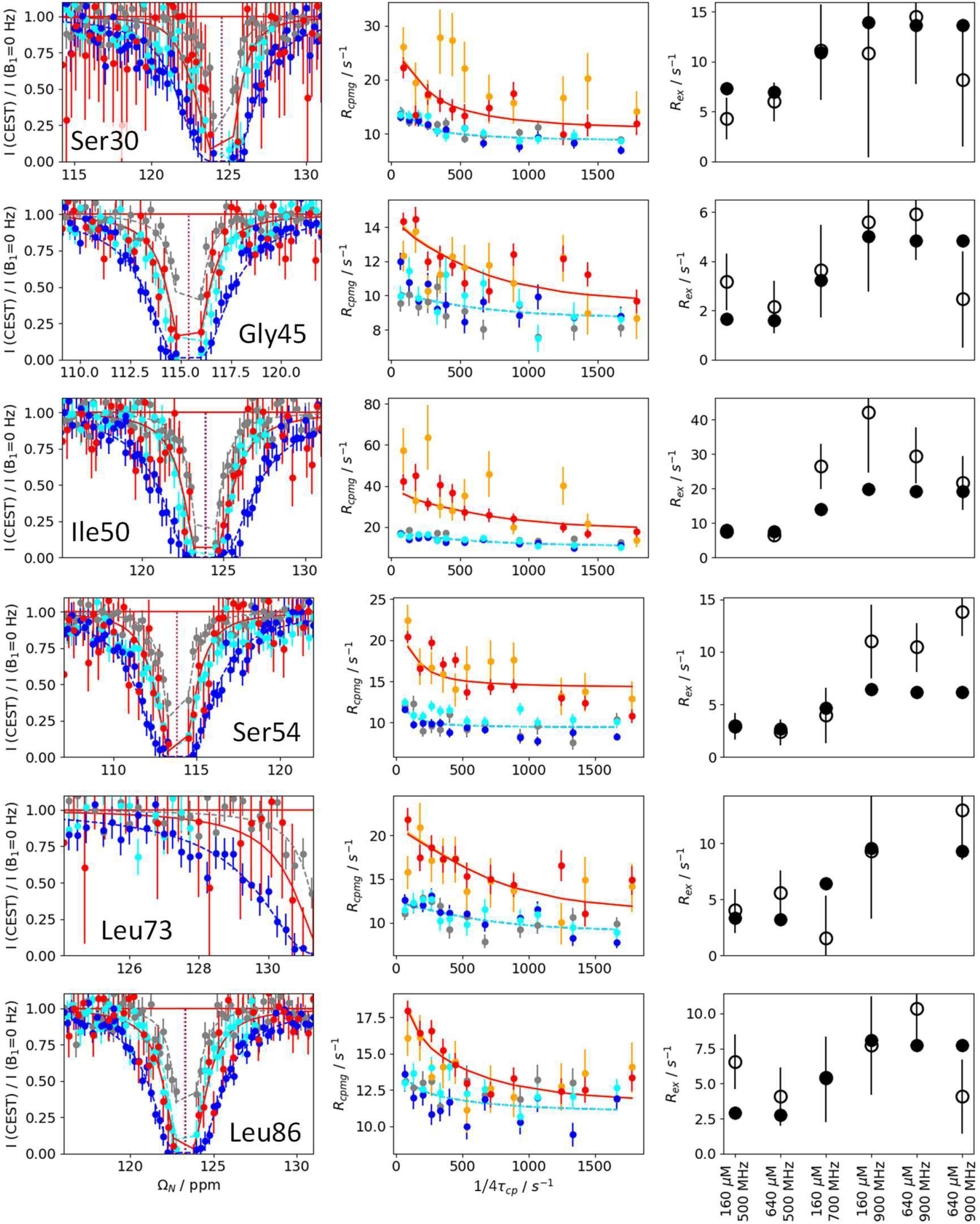
^15^N-relaxation dispersion reveals three-site exchange. Six more residues included in the global fit are shown, expanding Fig. 5. Left panels: 15N-CEST at c = 640 µM, B_0_ = 800 MHz, B_1_ = 35 Hz (red) or at c = 160 µM and B_0_=700 MHz (grey – B_1_ = 15 Hz, cyan – B_1_ = 15 Hz, blue – B_1_ = 50 Hz); center panels: ^15^N-CPMG at c = 160 µM (grey, orange), c = 640 µM (blue, cyan, red), B_0_ = 500 MHz (grey, cyan, blue), B_0_ = 900 MHz (orange, red); right panels: ^15^N-high power spin-lock experiments for *R*_*ex*_ determination, with conditions listed along the x axis with experimental (open circles) and calculated (filled circles) values. For the global fit, ^15^N-CEST data were converted to *R*_1ρ_-type data as described elsewhere (Palmer and Koss, 2019), using *R*_1ρ_ multi-site approximations for initial calculations (Koss et al., 2017). ^15^N-CPMG data were fitted with an the exact solution or an approximation (see STAR Methods) for the triangular state (Koss et al., 2018).

**Fig. S10:**
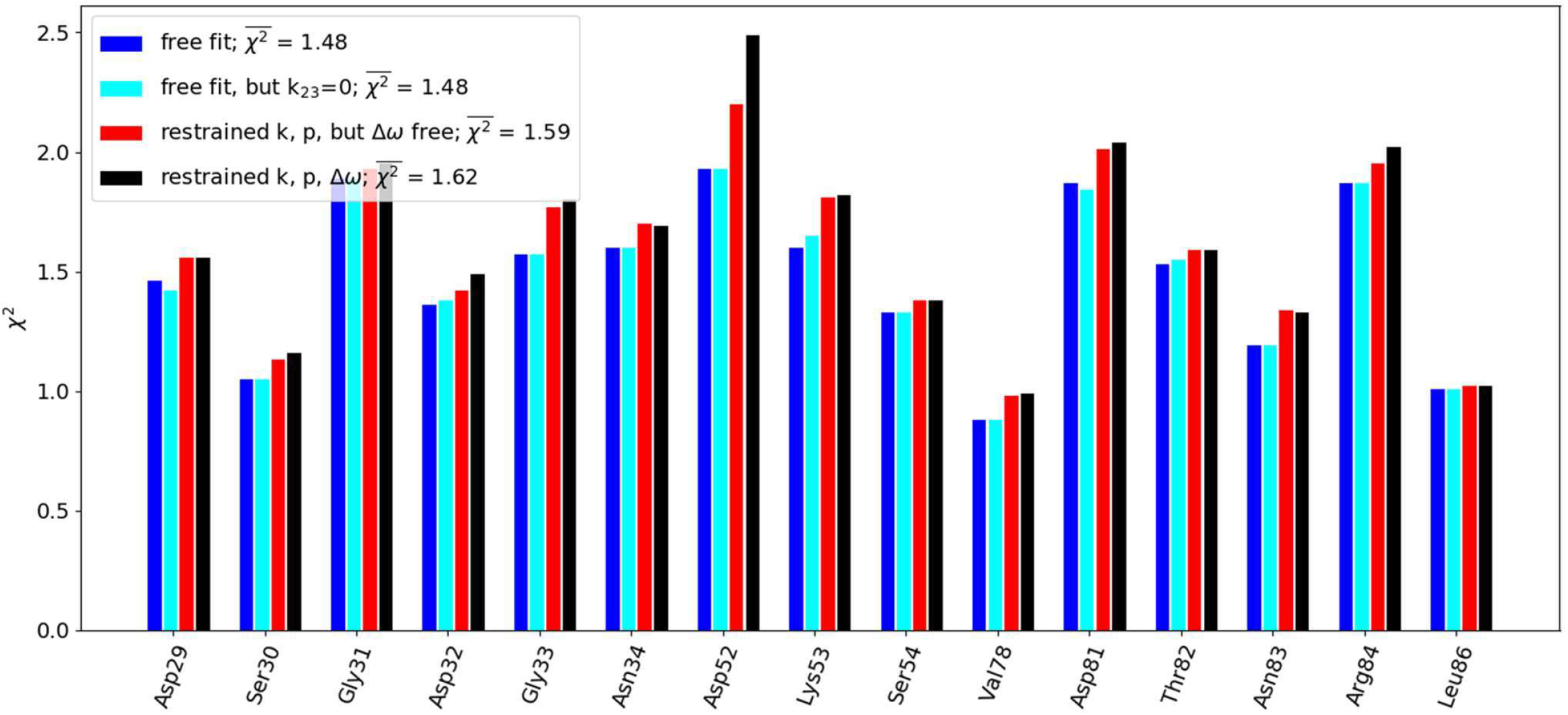
Validation of minor state chemical shifts. ^15^N-CPMG relaxation dispersion data from BC, DE and FG loop residues were used for this analysis. Fits for ^15^N relaxation dispersion data (see also Fig. S9) were performed for each residue individually rather than globally, even with flat or poor relaxation dispersion profiles. This analysis aims to test whether our assumptions about sparsely populated state shifts are valid. Δω: Boundary conditions for new residues were generated as in the global fit: We used all known W2A and W2AW4A chemical shifts for setting Δω restraints. The results from the global fit (Fig. 5, Fig. S9) were used at starting parameters, if available, but boundary conditions were varied: (1) all parameters including k_23_+k_32_ free; (2) all parameters free but k_23_+k_32_ = 0; (3) only Δω free; (4) all parameters maximally restrained. If starting parameters were not available, they were generated from the boundary conditions. Leaving Δω unrestrained does not improve χ^2^. Leaving k_12_+k_21_, k_13_+k_31_, p_b_ and p_c_ free for each residue gives only very slightly lower χ^2^ than restraining it globally. Allowing chemical exchange between the minor sites B and C does not change χ^2^.

**Fig. S11:**
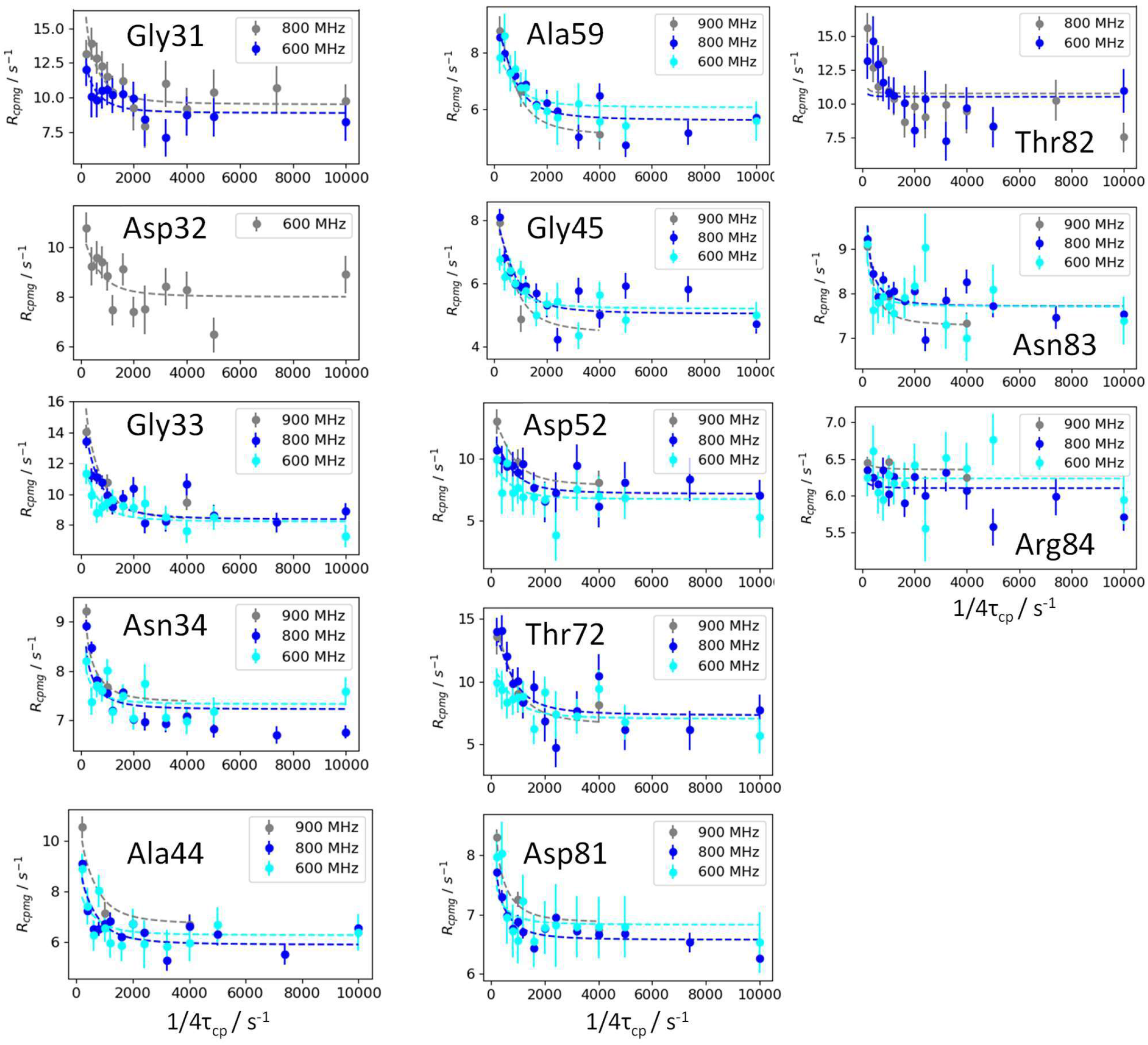
^1^H-relaxation of a deuterated Cad11-WT sample qualitatively corroborates the results from the ^15^N relaxation analysis. The ^1^H-CPMG curves, recorded at three different fields (600, 800, 900 MHz, see panel legends), are shown for residues for which shift predictions for Δ *ω*_AB_ and Δ *ω*_AB_ were available. The data are only of moderate quality, mostly due to degradation of the deuterated samples. These data serve to qualitatively corroborate the characterization of the sparsely populated states. All global fits used the rate constants and populations generated from the ^15^N fits as starting parameters and converged reliably, with resulting parameters which are neither reliable nor quantitatively comparable to the main results due to concentration and temperature discrepancies. The figure shows the result of a global fit which uses all predicted Δ *ω*_AB_ and Δ *ω*_AB_ from mutant chemical shifts: δ H_c_ is δ H(W2A,folded), and δH_B_ is usually δ H(W2AW4A,folded), or a range between δ H(W2AW4A,folded) and δ H(W2AW4A,unfolded). The fit is poor for residue Thr82, even when boundaries for kinetic parameters are relaxed. With freely set Δ *ω*_AB_ and Δ *ω*_AC_, the Δ *ω*. for Thr82 becomes very large and global χ^2^ for the global fit is reduced from 1.71 (this figure) to 1.47. We suspect that either data quality for Thr82 is insufficient and/or an additional exchange process captured specifically for this residue, probably due to the proximity of its proton shift to the random coil region.

**Fig. S12:**
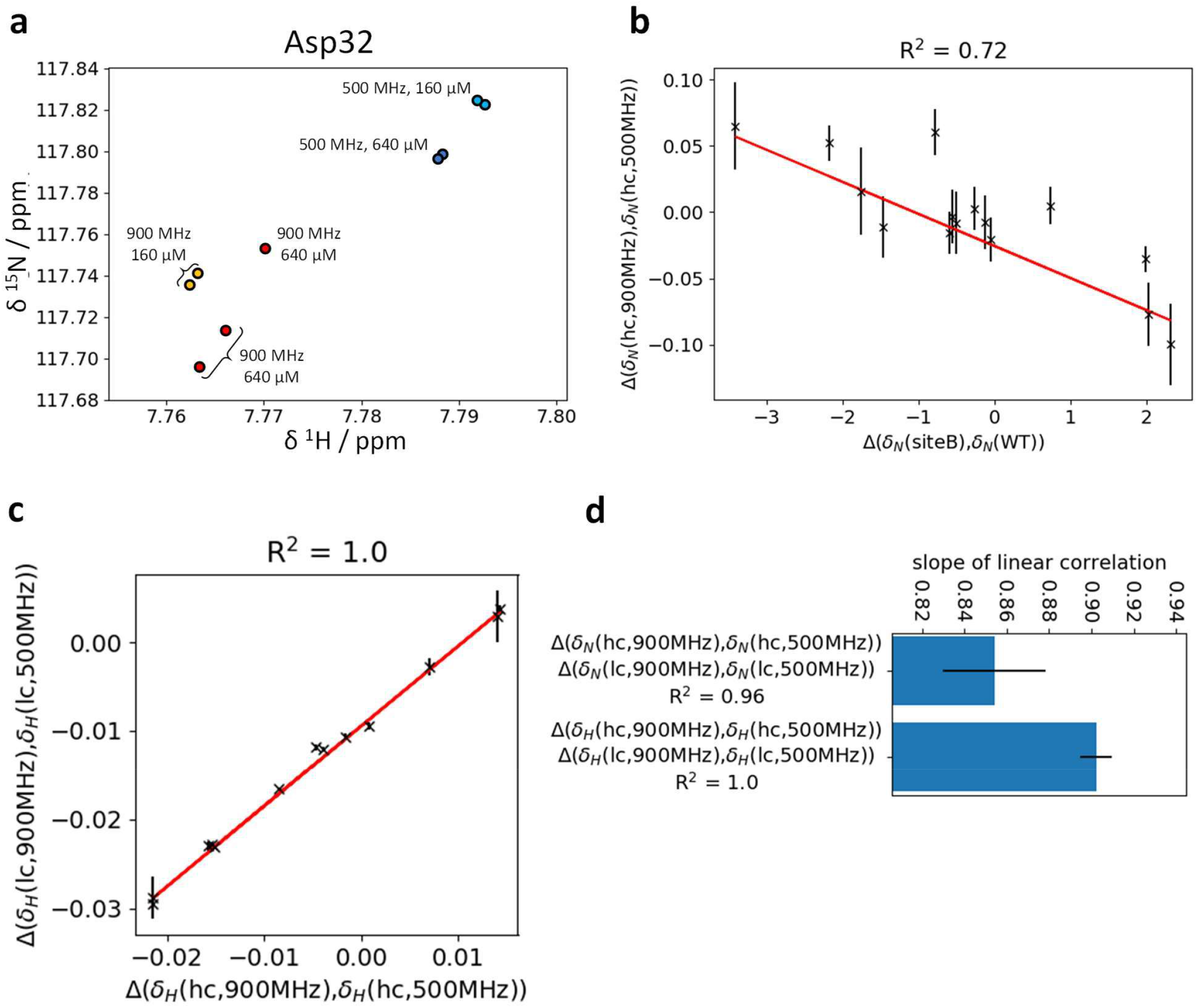
Field-dependent exchange-induced chemical shift perturbation (EI-CSPs) at different concentrations. **a:** For residue Asp32, ^1^H, ^15^N-TROSY-HSQC peak positions at 285K for different concentrations (160 µM and high 640 µM) and fields (500 and 900 MHz) are indicated. **b:** Correlation between Δδ_B0_ and chemical shift differences to identify the underlying exchange process. bΔδ_B0_ (here at 640 µM) show partial anticorrelation with the chemical shift difference between the two fast-intermediate exchanging states (minor state: fully strand exposed / W2AW4A-like). This correlation implies that the exchange process contributes in part to t.6so. **c and d:** Virtually perfect correlations between field-dependent chemical shifts at low vs. high concentration corroborates the presence and small errors for Δδ_B0_. The correlation plot is shown in panel c for ^1^H shifts. The fitted equation is Δδ_B0,1ow_ = (0.9*Δδ_H_^500-900^-0.009) ppm, implying that not only Δδ_B0_, but also the underlying exchange process is concentration dependent. The fitted slopes for the relationship between the Δδ_B0_ at low and high concentration for ^1^H and ^15^N nuclei are shown in panel d. The fact that the slopes are ≪ 1 therefore indicates that the underlying fast-intermediate exchange process with the fully A-strand-exposed state is concentration-dependent.

**Fig. S13:**
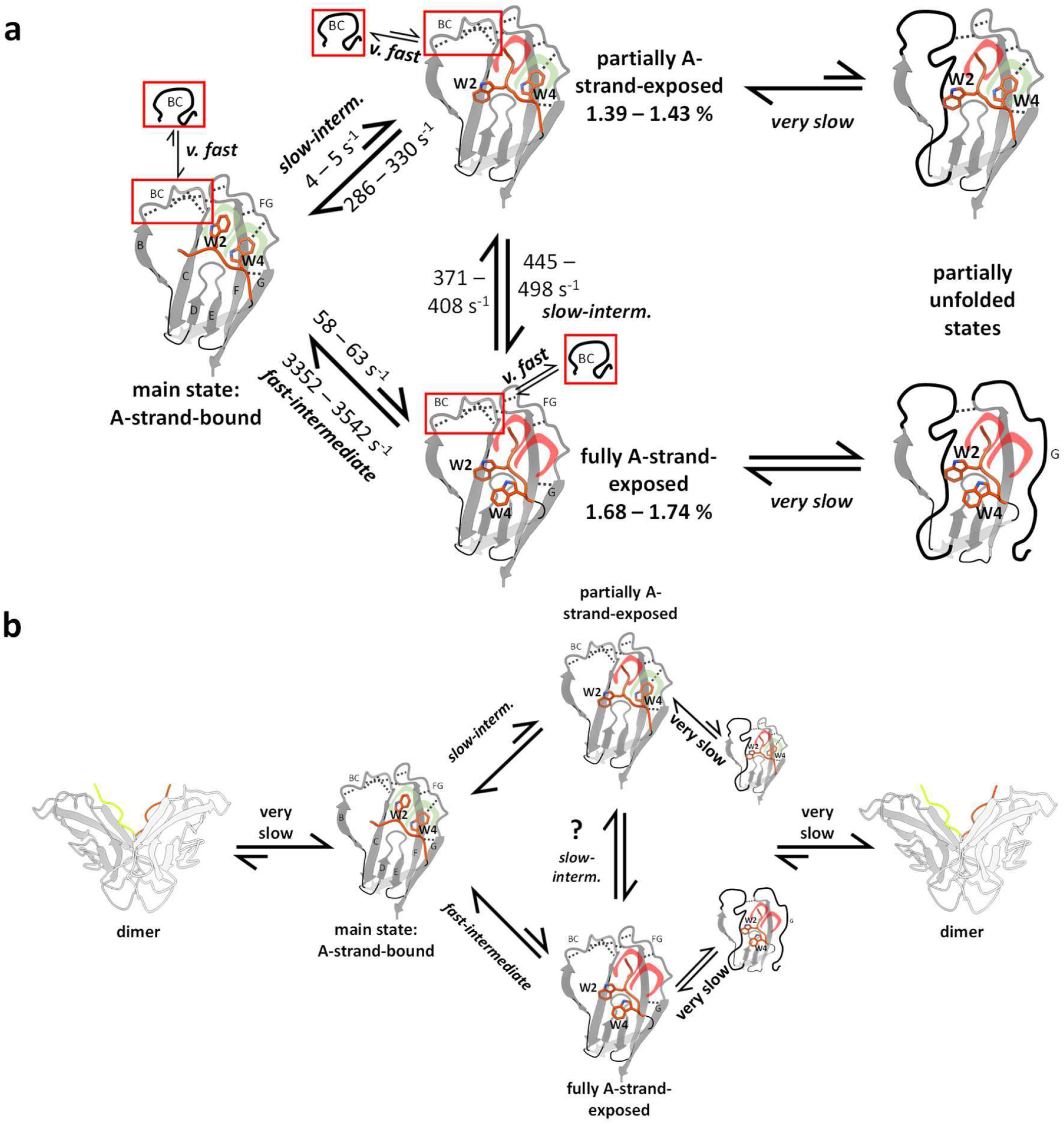
Topological context of strand-swapping equilibria. a. Triangular fitting results with exchange between sparsely populated sites. An alternative, triangular model fits to the same states as the linear “BAC” models, with different kinetic parameters. The local very fast equilibria for BC loop are shown in red insets, with the equilibrium arrow indicating general propensity towards the ordered or the RC state. Exemplary contacts in the locally ordered states are shown as dotted lines, RC sections are shown as black solid lines. The binding status of the Trp2 and Trp4 pockets are illustrated in red (not bound) and green (bound). Important strand and loop labels are indicated in the schematic, simplified models. Resulting rate constant and population ranges for c = 640 µM are shown; population data do not include partially unfolded states. For details see Table S3. **b. Competing dimerization pathways**. The fact that concentration dependency appears likely based on RD experiments and has been observed in the analysis of Δδ_B0_ implies that the exchange equilibrium discovered with RD experiments is on-pathway in context of more than one dimerization process: At least the one of the A-strand-exposed state is on-pathway, and at least one other competing pathway exists. This scheme provides an example for a kinetic Cadherin-11 dimerization scheme with two competing pathways, illustrating potential involvement of partially unfolded state(s) in one of the dimerization processes.

**Table S1:** ^1^H, ^15^N chemical shifts of folded (non-random-coil) states for various constructs under various conditions. All chemical shifts are listed in ppm.

| residue | W2A-Cad11-EC1 |  | W4A-Cad11-EC1 |  | W2AW4A-Cad11-EC1 |  | WT-Cad11-EC1 (160 μM), <sup>15</sup>N-TROSY CPMG experiments |  |  |  |  |  |  |  |
| --- | --- | --- | --- | --- | --- | --- | --- | --- | --- | --- | --- | --- | --- | --- |
|  | 800 MHz, 285 K, <sup>15</sup>N-TROSY-HSQC experiments |  |  |  |  |  | 500 MHz, 285K |  | 500 MHz, 298K |  | 900 MHz, 285K |  | 900 MHz, 298K |  |
|  | δH | δN | δH | δN | δH | δN | δH | δN | δH | δN | δH | δN | δH | δN |
| Thr14 | 8.16 | 116.7 |  |  | 8.16 | 116.8 | 8.162 | 116.53 | 8.14 | 116.43 | 8.158 | 116.54 | 8.136 | 116.42 |
| Gly15 |  |  |  |  | 6.81 | 110.1 |  |  |  |  |  |  |  |  |
| Leu20 |  |  |  |  |  |  | 8.567 | 128.89 |  |  | 8.551 | 128.89 |  |  |
| Asp29 | 7.42 | 121.4 | 7.28 | 120.8 | 7.51 | 121.9 | 7.332 | 120.86 | 7.379 | 120.94 | 7.309 | 120.82 | 7.38 | 120.92 |
| Ser30 | 8.75 | 122.5 |  |  | 8.75 | 122.3 | 9.045 | 124.49 | 9.012 | 124.31 | 9.051 | 124.51 | 9.015 | 124.32 |
| Gly31 | 9.02 | 111.3 | 9.20 | 111.5 | 8.99 | 111.5 | 9.277 | 111.51 | 9.287 | 111.53 | 9.267 | 111.48 | 9.292 | 111.51 |
| Asp32 | 7.78 | 118.2 | 7.78 | 117.9 | 7.85 | 118.5 | 7.793 | 117.82 | 7.846 | 118.01 | 7.762 | 117.73 | 7.847 | 117.99 |
| Gly33 | 8.83 | 109.4 | 8.96 | 109.3 | 8.69 | 109.3 | 9.051 | 109.39 | 9.063 | 109.38 | 9.038 | 109.36 | 9.069 | 109.35 |
| Asn34 | 8.36 | 118.2 | 8.33 | 118.2 | 8.36 | 118.2 | 8.371 | 118.3 | 8.414 | 118.31 | 8.348 | 118.27 | 8.415 | 118.3 |
| Ile35 |  |  |  |  |  |  | 7.184 | 118.39 | 7.247 | 118.57 | 7.134 | 118.3 | 7.243 | 118.55 |
| Lys36 | 8.97 | 129.1 | 9.03 | 130.1 | 8.98 | 129.8 | 9.077 | 130.13 | 9.095 | 130.01 | 9.058 | 130.14 | 9.094 | 130 |
| Tyr37 |  |  |  |  |  |  | 8.974 | 127.42 | 8.976 | 127.38 | 8.964 | 127.4 | 8.977 | 127.38 |
| Ile38 |  |  | 8.57 | 121.7 |  |  | 8.63 | 121.73 | 8.662 | 121.58 | 8.604 | 121.76 | 8.66 | 121.59 |
| Gly41 | 8.38 | 106.7 | 8.51 | 107.3 | 8.50 | 107.5 | 8.485 | 107.3 | 8.473 | 107.44 | 8.48 | 107.22 | 8.473 | 107.43 |
| Glu42 | 8.20 | 121.6 |  |  | 8.21 | 121.8 | 8.189 | 121.5 | 8.186 | 121.57 | 8.212 | 121.68 | 8.184 | 121.57 |
| Gly43 | 8.55 | 110.1 | 8.55 | 109.5 | 8.58 | 109.6 | 8.556 | 109.86 | 8.549 | 109.83 | 8.55 | 109.83 | 8.55 | 109.81 |
| Ala44 | 8.05 | 126.2 |  |  | 8.03 | 126.0 | 8.151 | 126.74 | 8.168 | 126.78 | 8.136 | 126.7 | 8.173 | 126.79 |
| Gly45 | 9.55 | 115.3 | 9.56 | 115.1 | 9.61 | 115.0 | 9.616 | 115.43 | 9.63 | 115.44 | 9.605 | 115.4 | 9.634 | 115.42 |
| Thr46 | 7.26 | 110.2 | 7.24 | 110.2 | 7.29 | 110.2 | 7.277 | 110.3 | 7.318 | 110.44 | 7.255 | 110.23 | 7.319 | 110.43 |
| Phe48 |  |  |  |  | 7.63 | 114.8 | 7.656 | 114.96 | 7.692 | 115.14 | 7.637 | 114.89 | 7.689 | 115.13 |
| Val49 |  |  |  |  |  |  | 8.892 | 119.38 | 8.935 | 119.53 | 8.865 | 119.3 | 8.937 | 119.51 |
| Ile50 |  |  |  |  |  |  | 8.803 | 123.91 | 8.783 | 123.97 | 8.799 | 123.85 | 8.781 | 123.96 |
| Asp51 |  |  |  |  |  |  | 8.454 | 127.12 |  |  | 8.451 | 127.18 |  |  |
| Asp52 | 9.03 | 126.1 |  |  | 8.97 | 126.1 | 9.075 | 126.34 | 9.04 | 126.39 | 9.079 | 126.3 | 9.042 | 126.37 |
| Lys53 | 8.23 | 118.7 | 8.27 | 119.0 |  |  | 8.315 | 119.34 | 8.322 | 119.16 | 8.299 | 119.36 | 8.319 | 119.14 |
| Ser54 |  |  | 8.18 | 113.7 |  |  | 8.202 | 113.77 | 8.186 | 113.62 | 8.198 | 113.79 | 8.186 | 113.61 |
| Gly55 | 8.48 | 110.8 | 8.44 | 110.5 | 8.43 | 110.1 | 8.537 | 110.19 | 8.576 | 110.38 | 8.517 | 110.09 | 8.578 | 110.36 |
| Asn56 |  |  |  |  |  |  | 7.924 | 118.7 | 7.933 | 118.78 | 7.912 | 118.63 | 7.937 | 118.76 |
| Ala59 |  |  |  |  |  |  | 9.144 | 123.25 | 9.175 | 123.32 | 9.125 | 123.2 | 9.175 | 123.32 |
| Thr62 |  |  |  |  | 7.91 | 115.2 | 7.991 | 115.35 | 7.974 | 115.08 | 7.986 | 115.39 | 7.973 | 115.06 |
| Glu66 |  |  |  |  |  |  | 8.225 | 117.41 | 8.244 | 117.43 | 8.207 | 117.38 | 8.245 | 117.43 |
| Ala69 |  |  |  |  |  |  | 8.402 | 124.59 | 8.398 | 124.51 | 8.395 | 124.58 | 8.399 | 124.49 |
| Tyr71 |  |  |  |  |  |  | 8.756 | 119.85 | 8.787 | 120.05 | 8.729 | 119.79 | 8.786 | 120.04 |
| Thr72 | 8.75 | 119.9 | 8.96 | 119.8 | 8.94 | 119.5 | 8.821 | 119.65 | 8.843 | 119.69 | 8.804 | 119.6 | 8.844 | 119.67 |
| Leu73 |  |  |  |  |  |  | 8.401 | 131.81 | 8.423 | 131.59 | 8.383 | 131.86 | 8.422 | 131.57 |
| Ala77 |  |  |  |  |  |  | 8.987 | 123.96 | 8.932 | 124.04 | 8.997 | 123.9 | 8.934 | 124.01 |
| Val78 | 8.85 | 117.2 |  |  | 8.91 | 117.3 | 8.645 | 115.14 | 8.701 | 115.45 | 8.614 | 115.03 | 8.698 | 115.43 |
| Asp79 |  |  |  |  |  |  | 8.543 | 122.67 | 8.565 | 122.79 | 8.528 | 122.62 | 8.568 | 122.78 |
| Arg80 | 8.38 | 128.1 |  |  | 8.42 | 128.2 | 8.646 | 128.57 |  |  | 8.649 | 128.51 |  |  |
| Asp81 | 8.51 | 115.6 | 8.55 | 115.8 | 8.49 | 115.5 | 8.615 | 116 | 8.619 | 116.1 | 8.604 | 115.95 | 8.62 | 116.09 |
| Thr82 | 7.83 | 106.6 | 7.83 | 106.7 | 7.86 | 106.6 | 7.908 | 106.91 | 7.941 | 106.94 | 7.885 | 106.88 | 7.937 | 106.93 |
| Asn83 | 8.30 | 117.3 | 8.40 | 117.6 | 8.32 | 117.3 | 8.492 | 117.76 | 8.51 | 117.78 | 8.48 | 117.72 | 8.513 | 117.76 |
| Arg84 | 7.50 | 120.7 | 7.49 | 120.6 | 7.50 | 120.7 | 7.526 | 120.47 | 7.553 | 120.41 | 7.509 | 120.46 | 7.554 | 120.39 |
| Leu86 |  |  |  |  |  |  | 8.853 | 123.3 | 8.842 | 123.31 | 8.849 | 123.29 | 8.843 | 123.32 |
| Ser90 |  |  |  |  |  |  | 8.344 | 116.2 | 8.314 | 116.1 | 8.345 | 116.22 | 8.315 | 116.07 |

**Table S2:** ^1^H, ^15^N chemical shifts and intensities of partially unfolded states/ RC peaks in Trp mutant constructs. The spectra which have been used to record these data have been recorded at 285 K and 800 MHz, of the TROSY-^15^N-HSQC-type. Intensities were normalized to the average of each construct’s RC intensities.

| residue | W2A-Cad11-EC1 |  |  | W4A-Cad11-EC1 |  |  | W2AW4A-Cad11-EC1 |  |  |
| --- | --- | --- | --- | --- | --- | --- | --- | --- | --- |
| | $\delta H$ / ppm | $\delta N$ / ppm | intensity | $\delta H$ / ppm | $\delta N$ / ppm | intensity | $\delta H$ / ppm | $\delta N$ / ppm | intensity |
| Thr14 | 8.08 | 117.9 | 0.53 | 8.08 | 118.0 | 0.19 | 8.08 | 118.1 | 0.54 |
| Gly22 | 8.29 | 111.8 | 0.28 | 8.29 | 111.7 | 0.13 | 8.30 | 111.9 | 0.31 |
| Ser30 | 8.24 | 117.3 | 0.82 | 8.24 | 117.3 | 0.60 | 8.24 | 117.4 | 0.82 |
| Gly31 | 8.39 | 110.9 | 1.30 | 8.40 | 110.9 | 0.83 | 8.40 | 110.9 | 1.02 |
| Asp32 | 8.13 | 120.2 | 2.37 | 8.13 | 120.2 | 1.02 | 8.13 | 120.2 | 1.71 |
| Gly33 | 8.39 | 109.3 | 1.19 | 8.40 | 109.3 | 0.63 | 8.40 | 109.3 | 0.91 |
| Asn34 | 8.20 | 118.6 | 1.16 | 8.20 | 118.6 | 0.49 | 8.20 | 118.5 | 0.76 |
| Gly41 | 8.34 | 110.7 | 0.64 | 8.34 | 110.7 | 0.25 | 8.34 | 110.7 | 0.36 |
| Glu42 | 8.24 | 120.4 | 0.80 | 8.24 | 120.4 | 0.30 | 8.24 | 120.4 | 0.53 |
| Gly43 | 8.45 | 110.1 | 0.64 | 8.45 | 110.2 | 0.64 | 8.46 | 110.1 | 0.41 |
| Ala44 | 8.08 | 124.0 | 0.41 | 8.10 | 124.0 | 0.21 | 8.09 | 124.0 | 0.26 |
| Gly45 | 8.34 | 108.1 | 0.40 | 8.34 | 108.1 | 0.26 | 8.34 | 108.0 | 0.29 |
| Ser54 | 8.21 | 116.0 | 0.57 | 8.20 | 115.7 | 0.24 | 8.19 | 115.7 | 0.36 |
| Gly55 | 8.28 | 110.4 | 0.60 | 8.26 | 110.4 | 0.37 | 8.27 | 110.5 | 0.52 |
| Thr62 | 7.86 | 114.4 | 0.57 | 7.86 | 114.4 | 0.14 | 7.86 | 114.5 | 0.22 |
| Gln76 |  |  | 0.00 |  |  | 0.00 | 8.32 | 120.0 | 0.24 |
| Ala77 | 7.98 | 118.8 | 0.37 | 7.99 | 118.9 | 0.28 | 7.99 | 118.9 | 0.76 |
| Val78 | 8.22 | 121.6 | 2.59 | 8.21 | 121.9 | 1.00 | 8.20 | 121.8 | 0.77 |
| Asp81 | 8.33 | 120.8 | 1.03 | 8.34 | 120.8 | 0.76 | 8.34 | 120.8 | 1.14 |
| Thr82 | 7.96 | 114.1 | 0.91 | 7.97 | 114.1 | 1.54 | 7.96 | 114.1 | 1.41 |
| Asn83 | 8.34 | 120.5 | 0.81 | 8.35 | 120.6 | 1.53 | 8.34 | 120.5 | 1.16 |
| Arg84 | 7.96 | 122.1 | 0.95 | 7.97 | 122.1 | 1.76 | 7.96 | 122.1 | 1.54 |
| Ser90 | 8.34 | 116.3 | 1.13 | 8.34 | 116.3 | 1.97 | 8.34 | 116.3 | 1.60 |
| Phe92 | 8.11 | 121.1 | 1.08 | 8.11 | 121.0 | 1.57 | 8.11 | 121.1 | 1.54 |
| Ile93 | 7.87 | 124.0 | 0.79 | 7.87 | 124.0 | 1.21 | 7.87 | 124.0 | 1.23 |
| Val94 | 8.16 | 126.2 | 0.97 | 8.16 | 126.3 | 1.52 | 8.16 | 126.3 | 1.45 |
| Lys95 | 8.33 | 127.0 | 1.16 | 8.33 | 127.0 | 1.92 | 8.33 | 127.1 | 1.82 |
| Gln97 | 8.38 | 125.1 | 1.59 | 8.39 | 125.1 | 3.07 | 8.39 | 125.1 | 2.30 |
| Asp98 | 8.00 | 128.6 | 2.35 | 8.00 | 128.5 | 3.59 | 8.00 | 128.6 | 3.04 |

**Table S3:** Kinetic parameter results. Parameters and their boundaries corresponding to the global fitting stages described in the methods text. The starting parameters for stages 1 and 2 were randomly selected from the parameter range defined by the boundaries. Stage 2 is being performed with and without additional restrictions for Δω and only 4 ambiguous Δω (see Table S4). The results from the stage 2 calculations with restrictions are very similar, and the smallest χ^2^ result set was chosen as starting parameter set for stage 3. The results for stage 3 are obtained as the average of the results from 16 runs with combinations of 4 ambiguous Δω. The results for stages 3 and 4 are shown for k_23_ + k_32_ = 0 or k_23_ + k_32_ > 0. The errors are then obtained by a bootstrapping process. Even though the parameter ranges for different concentrations overlap, we find that generally p_c_ (640 µM) < p_c_ (160 µM), k_12_+k_21_ (640 µM) > k_12_+k_21_ (160 µM) (if k_23_ + k_32_ = 0) and k_13_+k_31_ (640 µM) > k_13_+k_31_ (160 µM).

| parameter | unit | boundaries stage 1 |  | boundaries stage 2-4 |  | start stage 3 | results stage 3 | error range (stage 4) | par(640μM) / par(160mM) | results stage 3 | error range (stage 4) | par(640μM) / par(160mM) |
| --- | --- | --- | --- | --- | --- | --- | --- | --- | --- | --- | --- | --- |
|  |  | min | max | min | max |  |  |  |  |  |  |  |
| | | | | | | | | $k_{23} + k_{32} = 0$ | | | $k_{23} + k_{32} > 0$ | |
| $p_b$ (160 μM) | % | 0.5 | 50 | 0.5 | 10 | 1.59 | 1.57 | 1.54 - 1.59 | 0.99 - 1.00 | 1.72 | 1.70 - 1.75 | 0.98 - 1.00 |
| $p_b$ (640 μM) | % | 0.5 | 50 | 0.5 | 10 | 1.59 | 1.57 | 1.55 - 1.58 | | 1.70 | 1.68 - 1.74 | |
| $p_c$ (160 μM) | % | 0.5 | 50 | 0.5 | 10 | 1.64 | 1.63 | 1.62 - 1.63 | 0.96 - 0.97 | 1.48 | 1.45 - 1.49 | 0.93 - 0.97 |
| $p_c$ (640 μM) | % | 0.5 | 50 | 0.5 | 10 | 1.57 | 1.57 | 1.56 - 1.58 | | 1.41 | 1.39 - 1.43 | |
| $k_{12} + k_{21}$ (160 μM) | $s^{-1}$ | 1 | 10000 | 1000 | 8000 | 3109 | 3035 | 2930 - 3110 | 1.03 - 1.07 | 3427 | 3340 - 3539 | 0.99 - 1.04 |
| $k_{12} + k_{21}$ (640 μM) | $s^{-1}$ | 1 | 10000 | 1000 | 8000 | 3231 | 3179 | 3122 - 3217 | | 3485 | 3410 - 3606 | |
| $k_{13} + k_{31}$ (160 μM) | $s^{-1}$ | 100 | 900 | 100 | 900 | 663 | 669 | 664 - 674 | 1.02 - 1.06 | 296 | 266 - 316 | 1.05 - 1.16 |
| $k_{13} + k_{31}$ (640 μM) | $s^{-1}$ | 100 | 900 | 100 | 900 | 700 | 694 | 686 - 704 | | 325 | 290 - 334 | |
| $k_{23} + k_{32}$ (160 μM) | $s^{-1}$ | | | | | | | | | 857 | 860 - 900 | 0.96 - 1.05 |
| $k_{23} + k_{32}$ (640 μM) | $s^{-1}$ | | | | | | | | | 857 | 816 - 900 | |
| $\chi^2$ | | | | | | | 1.281 | 1.280 - 1.284 | | 1.272 | 1.268 - 1.275 | |

**Table S4:**
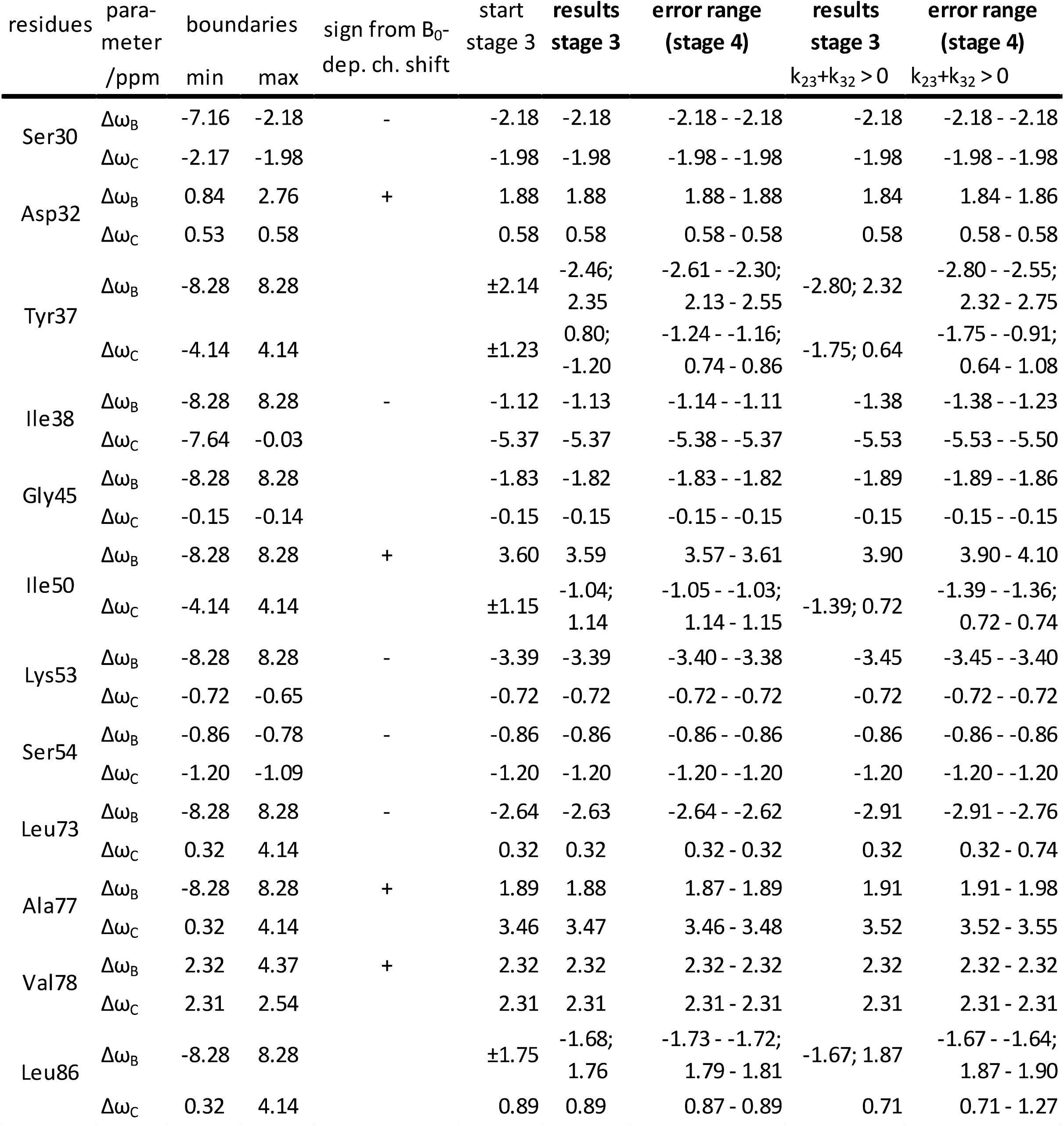
Chemical shift parameter results. Chemical shifts between the main state and the sparsely populated states B and C for each residue are listed. The boundaries apply to all stages. Starting parameters for stage 1 and 2 were randomly selected from the range defined by the boundaries. Field-dependent chemical shift perturbations indicate the sign for several Δω_B_, as indicated in the table. This restriction is used to define the starting parameters for stage 3. Also, Δω_B_(G45) is assumed to be negative (the chemical shift corresponding to the positive value would be unlikely). The results for stages 3 and 4 are shown for k_23_ + k_32_ = 0 or k_23_ + k_32_ > 0.

**Table S5:** *R*_*2,0*_ parameter results. *R*_*2,0*_ was in many cases expressed as combinations of different parameters to flexibly implement certain boundary conditions. The five sets are combined as follows: ^15^N-CPMG, 500 MHz: set 1; ^15^N-CPMG, 900 MHZ: set 1 * set 2; ^15^N-CEST 700 MHz: set 4 * set 5; ^15^N-CEST 800 MHz: set 1 * set 3. We assume *R*_*2,0*_ (^15^N-CPMG 900 MHz) *R*_*2,0*_ (^15^N-CPMG 500 MHz). The boundary condition *R*_*2,0*_ (^15^N-CEST 800 MHz) *R*_*2,0*_ (^15^N-CPMG 500 MHz) is debatable, for example due to the guessed uniform *R*_*1*_ contribution in ^15^N-CEST experiments. Removing this assumption leads to sometimes unreasonably small *R*_*2,0*_ (^15^N-CEST 800 MHz) with minimal changes of χ^2^ or other parameters. *R*_*ex*_ experiments do not require *R*_*2,0*_ for fitting which reduced any remnant bias introduced by assumptions about *R*_*2,0*_.

| boundaries | set 1 |  | set 2 |  | set 3 |  | set 4 |  | set 5 |  |
| --- | --- | --- | --- | --- | --- | --- | --- | --- | --- | --- |
| | $R_{2,0}$ (500 MHz, TROSY, CPMG) | | $R_{2,0}$ (900 MHz, TROSY, CPMG) / $R_{2,0}$ (500 MHz, TROSY) | | $R_{2,0}$ (800 MHz) / $R_{2,0}$ (500 MHz, TROSY, CEST) | | $R_{2,0}$ (500 MHz, non-TROSY) | | $R_{2,0}$ (700 MHz, non-TROSY, CEST) / $R_{2,0}$ (500 MHz, non-TROSY) | |
|  | 2 - 12 |  | 1 - 3 |  | 1 - 3 |  | 2-11 |  | 1 - 3 |  |
|  | av | err | av | err | av | err | av | err | av | err |
| Ser30 | 8.65 | 8.65 - 8.66 | 2.45 | 2.45 - 2.45 | 1.72 | 1.72 - 1.72 | 7.02 | 7.00 - 7.04 | 2.73 | 2.73 - 2.73 |
| Asp32 | 7.04 | 7.04 - 7.04 | 1.00 | 1.00 - 1.00 | 1.15 | 1.14 - 1.15 | 9.32 | 9.32 - 9.32 | 1.00 | 1.00 - 1.00 |
| Tyr37 | 5.86 | 5.85 - 5.87 | 1.28 | 1.28 - 1.29 | 1.10 | 1.10 - 1.11 | 9.40 | 9.40 - 9.40 | 1.22 | 1.22 - 1.22 |
| Ile38 | 9.26 | 9.25 - 9.26 | 2.34 | 2.34 - 2.35 | 2.21 | 2.20 - 2.21 | 7.98 | 7.98 - 7.99 | 1.83 | 1.83 - 1.83 |
| Gly45 | 9.21 | 9.13 - 9.31 | 1.00 | 1.00 - 1.00 | 1.00 | 1.00 - 1.00 | 9.10 | 9.09 - 9.11 | 1.49 | 1.49 - 1.49 |
| Ile50 | 6.86 | 6.74 - 7.05 | 1.27 | 1.26 - 1.27 | 1.77 | 1.77 - 1.78 | 6.80 | 6.79 - 6.81 | 1.00 | 1.00 - 1.00 |
| Lys53 | 9.52 | 9.52 - 9.52 | 2.07 | 2.07 - 2.07 | 2.30 | 2.30 - 2.30 | 9.21 | 9.20 - 9.22 | 2.83 | 2.81 - 2.83 |
| Ser54 | 9.04 | 9.03 - 9.05 | 1.00 | 1.00 - 1.00 | 1.60 | 1.60 - 1.60 | 8.26 | 8.25 - 8.26 | 1.29 | 1.24 - 1.34 |
| Leu73 | 8.68 | 8.67 - 8.68 | 1.24 | 1.23 - 1.24 | 1.46 | 1.46 - 1.47 | 8.94 | 8.93 - 8.95 | 1.17 | 1.16 - 1.20 |
| Ala77 | 10.34 | 10.34 - 10.34 | 2.41 | 2.41 - 2.41 | 2.79 | 2.79 - 2.79 | 9.35 | 9.34 - 9.35 | 2.30 | 2.30 - 2.31 |
| Val78 | 10.89 | 10.86 - 10.94 | 1.35 | 1.35 - 1.36 | 1.47 | 1.47 - 1.47 | 11.00 | 11.00 - 11.00 | 1.00 | 1.00 - 1.00 |
| Leu86 | 11.37 | 11.33 - 11.40 | 1.07 | 1.06 - 1.07 | 1.51 | 1.51 - 1.51 | 7.60 | 7.59 - 7.60 | 1.31 | 1.31 - 1.31 |

**Table S6:** Experimental conditions of ^15^N-R_ex_ experiments. SL: spin lock power; ns: number of scans. Related to STAR Methods, section ^15^N-R_ex_ experiments.

| B<sub>0</sub> / MHz | conc /<br>μM | T / ms | SL / kHz | ns |
| --- | --- | --- | --- | --- |
| 500 | 160 | 50 | 2.5 | 48 |
| 700 | 160 | 30 | 3.5 | 32 |
| 900 | 160 | 30 | 2.5 | 32 |
| 500 | 640 | 50 | 2.5 | 8 |
| 900 | 640 | 30 | 2.5 | 8 |
| 900 | 640 | 30 | 2.5 | 8 |

**Table S7:** Experimental conditions of ^15^N-CPMG experiments. ns: number of scans. Related to STAR Methods, section ^15^N-CPMG experiments.

| B<sub>0</sub> /<br>MHz | conc / μM | T / ms | ns |
| --- | --- | --- | --- |
| 500 | 2.475 | 60 | 40 |
| 900 | 2.475 | 45 | 32 |
| 500 | 9.9 | 30 | 8 |
| 500 | 9.9 | 30 | 8 |
| 900 | 9.9 | 45 | 8 |

## References

Brasch, J., Harrison, O. J., Honig, B. & Shapiro, L. 2012. Thinking outside the cell: how cadherins drive adhesion. Trends Cell Biol, 22, 299–310.

Brasch, J., Katsamba, P. S., Harrison, O. J., Ahlsen, G., Troyanovsky, R. B., Indra, I., Kaczynska, A., Kaeser, B., Troyanovsky, S., Honig, B. & Shapiro, L. 2018. Homophilic and Heterophilic Interactions of Type II Cadherins Identify Specificity Groups Underlying Cell-Adhesive Behavior. Cell Rep, 23, 1840–1852.

Byeon, I. J., Louis, J. M. & Gronenborn, A. M. 2004. A captured folding intermediate involved in dimerization and domain-swapping of GB1. J Mol Biol, 340, 615–25.

Chang, S. K., Gu, Z. & Brenner, M. B. 2010. Fibroblast-like synoviocytes in inflammatory arthritis pathology: the emerging role of cadherin-11. Immunol Rev, 233, 256–66.

Daoud, M., Cotton, J. P., Farnoux, B., Jannink, G., Sarma, G., Benoit, H., Duplessix, C., Picot, C. & De Gennes, P. G. 1975. Solutions of Flexible Polymers. Neutron Experiments and Interpretation. Macromolecules, 8, 804–818.

Delaglio, F., Grzesiek, S., Vuister, G. W., Zhu, G., Pfeifer, J. & Bax, A. 1995. NMRPi pe: A multidimensional spectral processing system based on UNIX pipes. J Biomol NMR, 6, 277–293.

Esposito, L. & Daggett, V. 2005. Insight into ribonuclease A domain swapping by molecular dynamics unfolding simulations. Biochemistry, 44, 3358–68.

Garcia De La Torre, J., Huertas, M. L. & Carrasco, B. 2000. HYDRONMR: prediction of NMR relaxation of globular proteins from atomic-level structures and hydrodynamic calculations. J Magn Reson, 147, 138–46.

Gronenborn, A. M. 2009. Protein acrobatics in pairs--dimerization via domain swapping. Curr Opin Struct Biol, 19, 39–49.

Harrison, O. J., Bahna, F., Katsamba, P. S., Jin, X., Brasch, J., Vendome, J., Ahlsen, G., Carroll, K. J., Price, S. R., Honig, B. & Shapiro, L. 2010. Two-step adhesive binding by classical cadherins. Nat Struct Mol Biol, 17, 348–57.

Igumenova, T. I. & Palmer, A. G. 2006. Off-resonance TROSY-selected R1rho experiment with improved sensitivity for medium-and high-molecular-weight proteins. J Am Chem Soc, 128, 8110–1.

Kang X., Zhong, N., Zou, P., Zhang, S., Jin, C. & Xia, B. 2012. Foldon unfolding mediates the interconversion between M(pro)-C monomer and 3D domain-swapped dimer. Proc Natl Acad Sci U S A, 109, 14900–5.

Koss, H., Rance, M. & Palmer, A. G. 2017. General expressions for R1rho relaxation for N-site chemical exchange and the special case of linear chains. J Magn Reson, 274, 36–45.

Koss, H., Rance, M. & Palmer, A. G. 2018. General Expressions for Carr-Purcell-Meiboom-Gill Relaxation Dispersion for N-Site Chemical Exchange. Biochemistry, 57, 4753–4763.

Lescop, E., Kern, T. &Brutscher, B. 2010. Guidelines for the use of band-selective radiofrequency pulses in hetero-nuclear NMR: Example of longitudinal-relaxation-enhanced BEST-type 1H–15N correlation experiments. J Magn Reson, 203, 190–198.

Li, Y., Altorelli, N. L., Bahna, F., Honig, B., Shapiro, L. & Palmer, A. G. 2013. Mechanism of E-cadherin dimerization probed by NMR relaxation dispersion. Proc Natl Acad Sci USA, 110, 16462–7.

Liu, L., Byeon, I. J., Bahar, I. & Gronenborn, A. M. 2012. Domain swapping proceeds via complete unfolding: a 19F-and 1H-NMR study of the Cyanovirin-N protein. J Am Chem Soc, 134, 4229–35.

Liu, Y., Gotte, G., Libonati, M. & Eisenberg, D. 2001. A domain-swapped RNase A dimer with implications for amyloid formation. Nat Struct Biol, 8, 211–4.

Liu, Z. & Huang, Y. 2013. Evidences for the unfolding mechanism of three-dimensional domain swapping. Protein Sci, 22, 280–6.

Miloushev, V. Z., Bahna, F., Ciatto, C., Ahlsen, G., Honig, B., Shapiro, L. & Palmer, A. G. 2008. Dynamic properties of a type II cadherin adhesive domain: implications for the mechanism of strand swapping of classical cadherins. Structure, 16, 1195–205.

Moschen, T. & Tollinger, M. 2014. A kinetic study of domain swapping of Protein L. Phys Chem Chem Phys, 16, 6383–90.

Murray, V., Huang, Y., Chen, J., Wang, J. & Li, Q. 2012. A novel bacterial expression method with optimized parameters for very high yield production of triple-labeled proteins. Methods Mol Biol, 831, 1–18.

Nguyen, T. T., Ghirlando, R., Roche, J. & Venditti, V. 2021. Structure elucidation of the elusive Enzyme I monomer reveals the molecular mechanisms linking oligomerization and enzymatic activity. Proc Natl Acad Sci USA, 118, e2100298118.

Palmer, A.G. & Koss, H. 2019. Chemical Exchange. Methods Enzymol, 615, 177–236.

Patel, S. D., Ciatto, C., Chen, C. P., Bahna, F., Rajebhosale, M., Arkus, N., Schieren, I., Jessell, T. M., Honig, B., Price, S. R. & Shapiro, L. 2006. Type II cadherin ectodomain structures : implications for classical cadherin specificity. Cell, 124, 1255–68.

Quinlan, R. J. & Reinhart, G. D. 2005. Baroresistant buffer mixtures for biochemical analyses. Anal Biochem, 341, 69–76.

Rousseau, F., Schymkowitz, J. W. & ltzhaki, L. S. 2003. The unfolding story of three-dimensional domain swapping. Structure, 11, 243–51.

Sfikakis, P. P., Vlachogiannis, N. I. & Christopoulos, P. F. 2017. Cadherin-11 as a therapeutic target in chronic, inflammatory rheumatic diseases. Clin Immunol, 176, 107–113.

Shapiro, L., Fannon, A. M., Kwong, P. D., Thompson, A., Lehmann, M. S., Grübel, G., Legrand, J.-F., Als Nielsen, J., Colman, D. R. & Hendrickson, W. A. 1995. Structural basis of cell-cell adhesion by cadherins. Nature, 374, 327–337.

Sivasankar, S., Zhang, Y., Nelson, W. J. & Chu, S. 2009. Characterizing the initial encounter complex in cadherin adhesion. Structure, 17, 1075–81.

Spadaccini, R., Ercole, C., Graziano, G., Wechselberger, R., Boelens, R. & Picone, D. 2014. Mechanism of 3D domain swapping in bovine seminal ribonuclease. FEBS J, 281, 842–50.

Vendome, J., Posy, S., Jin, X., Bahna, F., Ahlsen, G., Shapiro, L. & Honig, B. 2011. Molecular design principles underlying beta-strand swapping in the adhesive dimerization of cadherins. Nat Struct Mol Biol, 18, 693–700.

Vranken, W. F., Boucher, W., Stevens, T. J., Fogh, R. H., Pajon, A., Llinas, M., Ulrich, E. L., Markley, J. L., lonides, J. & Laue, E. D. 2005. The CCPN data model for NMR spectroscopy: development of a software pipeline. Proteins, 59, 687–96.

Yang, S., Cho, S. S., Levy, Y., Cheung, M. S., Levine, H., Wolynes, P. G. & Onuchic, J. N. 2004. Domain swapping is a consequence of minimal frustration. Proc Natl Acad Sci USA, 101, 13786–91.

Zhu, G., Xia, Y., Nicholson, L. K. & Sze, K. H. 2000. Protein dynamics measurements by TROSY-based NMR experiments. J Magn Reson, 143, 423–6.

